# Integration of Unpaired and Heterogeneous Clinical Flow Cytometry Data

**DOI:** 10.1101/2023.12.18.572157

**Authors:** Mike Phuycharoen, Verena Kaestele, Thomas Williams, Lijing Lin, Tracy Hussell, John Grainger, Magnus Rattray

## Abstract

We introduce the Unbiasing Variational Autoencoder (UVAE), a computational framework for the integration of unpaired biomedical data streams such as clinical flow cytometry. UVAE addresses batch effect correction and data alignment by training a semi-supervised model on partially labelled datasets, enabling simultaneous normalisation and integration of diverse data within a shared latent space. The framework implements a probabilistic model for batch effect normalisation and balances class contents during training to ensure accurate representation of underlying cell composition. We apply UVAE to integrate heterogeneous clinical flow cytometry data from COVID-19 patients. The integrated data enhances the statistical signal of cell types associated with disease severity, enables clustering of subpopulations without the impediment of batch effects, and improves the performance of longitudinal regression for predicting peak disease severity from temporal patient samples.

## Introduction

High-throughput technologies generate large-scale data, offering opportunities for identifying disease-associated biomarkers. Flow cytometry determines the relative abundance of proteins for each cell in a sample, enabling the identification of enriched cell types or shifts in activity that correlate with clinical outcomes. However, leveraging real-world clinical data is challenging due to the common use of multiple panels (unpaired data), changing feature sets, experimental variation (batch effects), and heterogeneous patient samples.

Methods for batch effect correction^1^ and unpaired data integration^2^ have been developed to address these issues. When feature sets are disparate, a simple approach is to eliminate unshared features, but this reduces available information. Alternatives include zero-padding missing values for neural network input^3^ or imputing them with regression^4^ or nearest-neighbour models^5^, though these methods are susceptible to batch effects. In cytometry, panels are often designed with shared markers to serve as a reference^5^. In such cases, one suitable integration solution is cyCombine^6^, which corrects batch effects within cell clusters using ComBat^7^ and merges panels by probabilistically drawing missing values from co-clustered cells. Such approaches require that shared markers are sufficient to link data streams.

Variational autoencoders (VAEs) offer advantages by performing corrections in a non-linear latent space^8^. Integration can be achieved by merging latent spaces through loss constraints, using either shared features^9^ or density-matching methods like Maximum Mean Discrepancy (MMD)^10,11^. VAE-based normalisation can be performed with latent arithmetic^12^ or conditional autoencoding^13^, though the latter may still capture batch-correlated information, which can be mitigated with additional loss terms like MMD^10^. A major challenge in all integration tasks is that unknown differences in class contents, such as varying cell-type proportions in clinical samples, can distort data alignment. Some methods address this by aligning data within known populations^6^ or by iteratively re-weighting the merging function to balance clusters^14^.

In this study, we introduce the Unbiasing Variational Autoencoder (UVAE) to integrate heterogeneous, unpaired datasets with changing feature sets. Our model separates confounding variability from a shared latent space, allowing the unification of datasets while performing simultaneous normalisation, merging, and class inference. A key component of our approach is the ability to account for differing class proportions between data series during integration, preventing over-alignment of biologically distinct samples. We demonstrate UVAE’s performance on synthetic data and apply it to a complex clinical flow cytometry dataset from COVID-19 patients, showing that the integrated data enhances biomarker discovery and improves downstream predictive modeling.

## Results

### UVAE framework integrates heterogeneous data through modular, constrained autoencoders

The development of our method was shaped by challenges encountered during the analysis of clinical blood samples from COVID-19 patients. Samples were processed using two main flow cytometry panel types: a *lineage* panel to identify immune cell lineages and a *chemokine* panel to quantify chemokine and cytokine levels. Over the 14-month data collection period, the markers used in these panels were altered due to changes in reagent availability and scientific focus, resulting in a dataset with substantial experimental variation, batch effects, and changing feature sets. This heterogeneity motivated the development of a method to integrate and harmonise the data for downstream analysis.

The UVAE is a deep learning framework designed to address these challenges. The overall workflow is illustrated in Figure 1. For each unique marker set, optionally partially labelled data is input to the UVAE algorithm, which performs simultaneous batch effect correction and panel integration. The model then generates a complete dataset with a uniform marker set and full class predictions for all cells. This homogeneous dataset is more conducive to downstream statistical evaluation and clustering, enhancing the power to identify cell types and markers correlated with disease severity.

**Figure 1.**
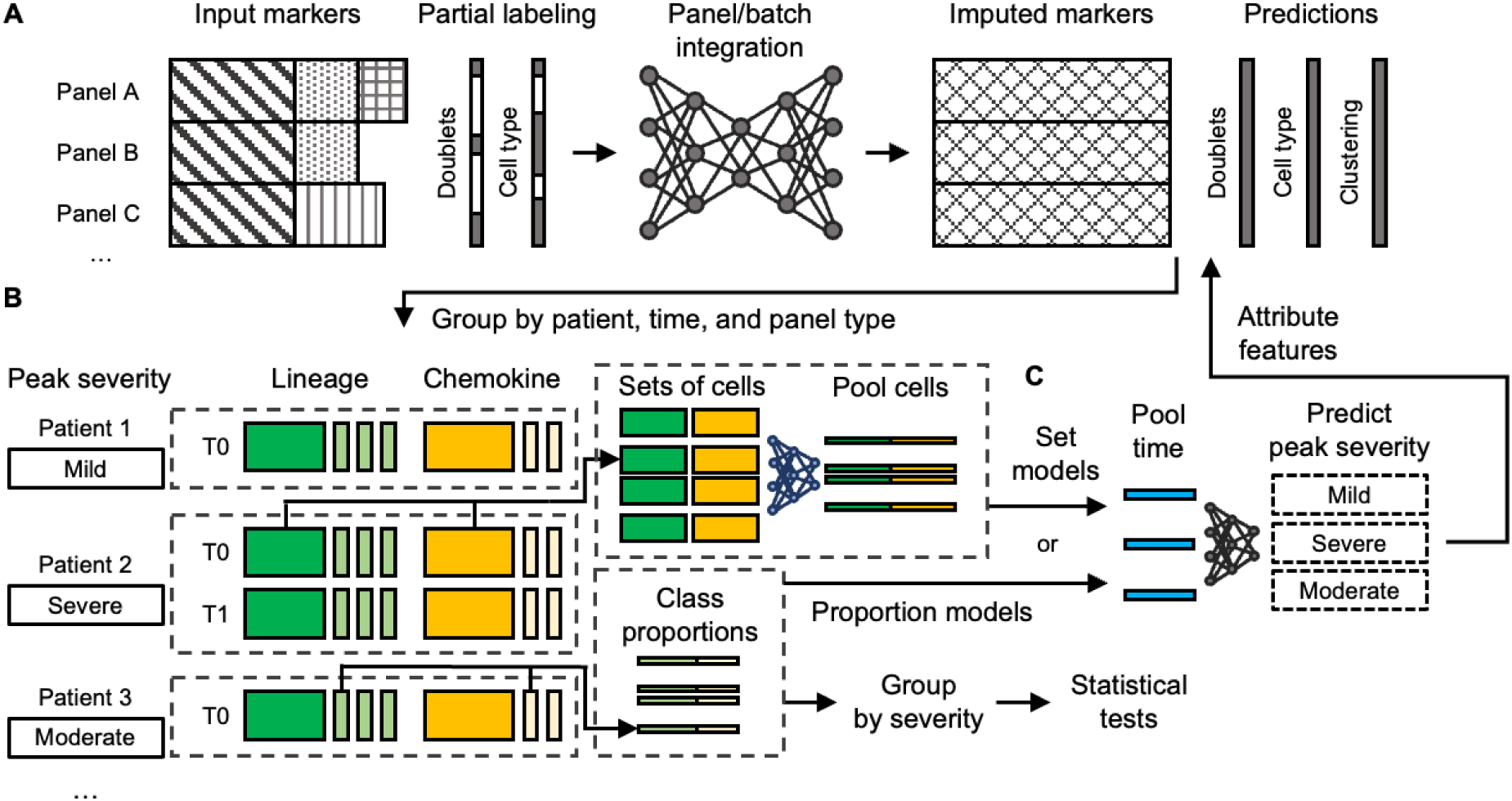
The UVAE analysis workflow. (A) Partially labelled flow cytometry data from multiple panels with partially shared markers is input to a deep learning algorithm performing batch effect correction and panel integration. The entire dataset is regenerated with a uniform marker set and full class predictions and can be optionally clustered. (B) Cell representations are split by panel type and grouped by patient-timepoint. Proportions of cell types, clusters, and corrected marker values are used to calculate enrichment by peak observed clinical severity. (C) Regression models predicting peak severity are trained using either class proportions or raw marker representations of cell sets as input, with a variable number of timepoints per patient. Gradient attribution is used to identify features contributing to an increase in predicted severity.

The core of the UVAE framework is a generative model constructed from multiple VAEs, one for each distinct input feature set (Figure 2). All VAEs project data into a shared-dimensionality latent space, where integration and normalisation are performed through a combination of user-specified constraints. These constraints are modules which define loss components that in combination influence data alignment.

**Figure 2.**
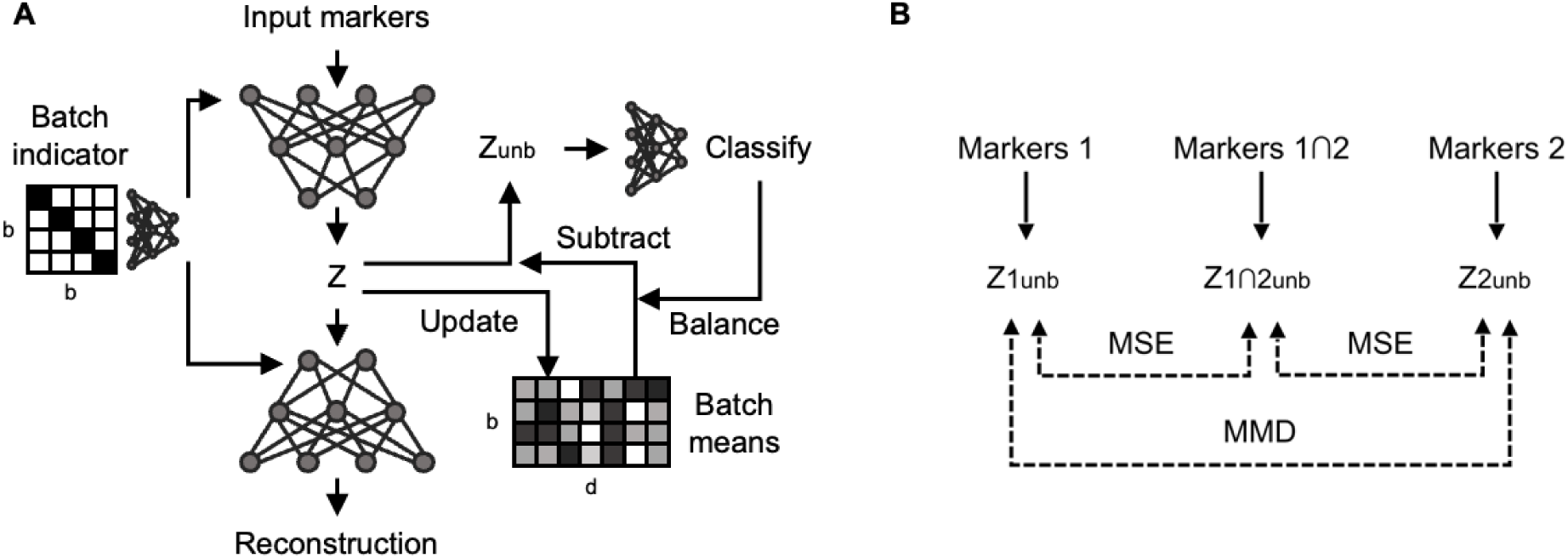
UVAE model architecture for batch effect correction and panel integration. (A) A single VAE is trained on one set of input markers, encoding data into a latent space (Z with dimensionality d) and decoding it back. Batch effect correction (for b batches) is performed by i) conditioning the encoder and decoder with a batch identifier and ii) offsetting the latent embedding to a common mean to produce an unbiased embedding (Zunb). Shared classifiers are trained on unbiased embeddings, and their predictions are used to balance the sample composition for calculating batch mean offsets. (B) Alignment of two input streams (Markers 1 and Markers 2) with partially overlapping features (Markers 1∩2). Separate VAEs are trained for each stream. An additional VAE is trained on the subset of shared features. Merging is performed by minimising the MSE loss between unbiased sample embeddings from each stream encoder and the shared encoder. Additional merging is performed by minimising an MMD loss calculated on class-balanced samples using the unbiased embeddings from the two stream encoders. Solid arrows indicate the encoding and unbiasing process within a single encoder. Dashed lines indicate merging losses applied between pairs of samples during training. Decoding and reconstruction loss is not shown for simplicity.

Batch effect correction is performed within each VAE using two concurrent mechanisms (Figure 2A). First, each VAE encoder and decoder can be conditioned with a batch identifier, which discourages the model from redundantly capturing batch-specific information. Second, latent arithmetic is used to align the embeddings. The mean of the latent embeddings for each batch is calculated using class-balanced samples, and individual sample embeddings are offset to a common reference mean. The resulting unbiased latent embeddings serve as input for shared classifiers and other integration constraints.

Integration of unpaired data streams is achieved by aligning the latent spaces of the different VAEs (Figure 2B). For data series that share a subset of markers, a *Subspace* constraint is applied. An additional VAE is trained on only the shared features, and the mean squared error (MSE) between its unbiased latent embeddings and those from the individual panel VAEs is minimised. To enable alignment without relying on shared features, a MMD loss is minimised between the unbiased latent embeddings of data from each series.

A key aspect of the UVAE is its ability to account for class imbalances between data series, which can distort distribution-matching approaches. The framework trains classifiers on available labels to predict cell types. These predictions are then used to balance the class proportions of samples used for calculating latent normalisation offsets and MMD loss, a process we refer to as resampling. This ensures that the aligned distributions represent proportional amounts of the underlying biological populations, preventing over-alignment.

This resampling can also be performed in an unsupervised manner by using cluster assignments derived independently for each batch. All model components are trained jointly via a common, non-adversarial optimisation objective, and hyper-parameters are selected via Bayesian optimisation to balance reconstruction accuracy with alignment performance.

### Systematic component validation on synthetic data demonstrates the benefits of integrated modeling

To quantitatively validate UVAE’s constituent components, we created a synthetic dataset with a known ground-truth alignment. A single, manually annotated flow cytometry sample was computationally split into three panels, each with three batches and distinct missing marker channels. To simulate biological heterogeneity, the underlying cell populations (defined by Gaussian Mixture Model [GMM] clustering) were distributed unevenly across batches and panels before batch-specific standardisation was applied, creating a dataset where simple normalisation cannot restore the correct alignment. Further details on data generation are available in the STAR Methods.

Using this dataset, we tested various combinations of UVAE’s batch correction and panel merging constraints. Model performance was evaluated using three metrics: imputation error (MSE) for the missing channels, batch mixing (iLISI), and ground-truth cluster separation (cLISI). Higher iLISI scores indicate better batch mixing, while lower cLISI scores indicate better preservation of biological structure. To account for class imbalances in the LISI calculation, scores were computed on subsets of data down-sampled to ensure equal class representation across batches (normalised LISI, see Figure S1 and STAR Methods). Hyper-parameters for each configuration were selected via Bayesian optimisation to balance reconstruction accuracy with these alignment scores. The specific components used in these models are detailed in Table 1.

**Table 1.**
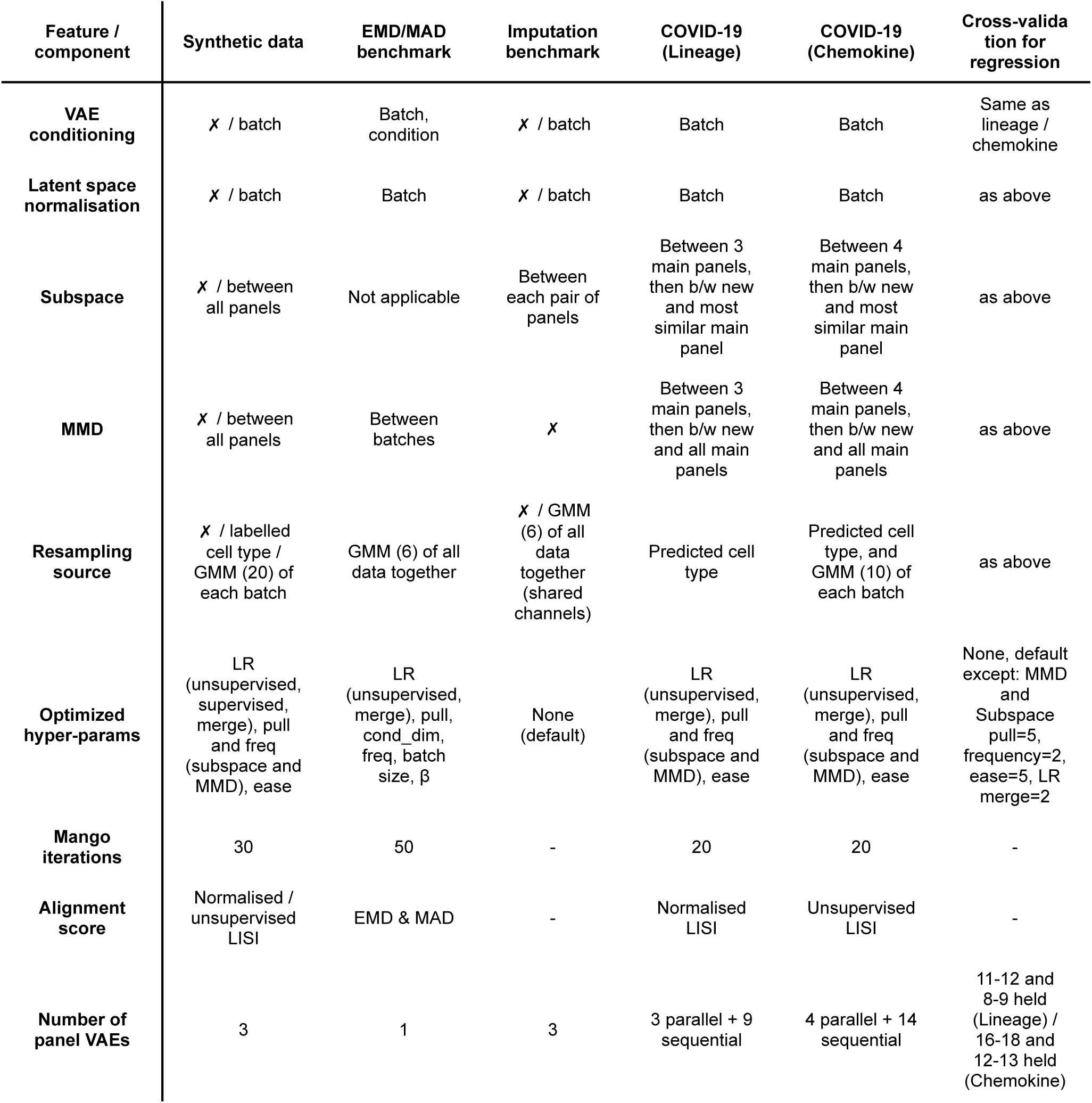
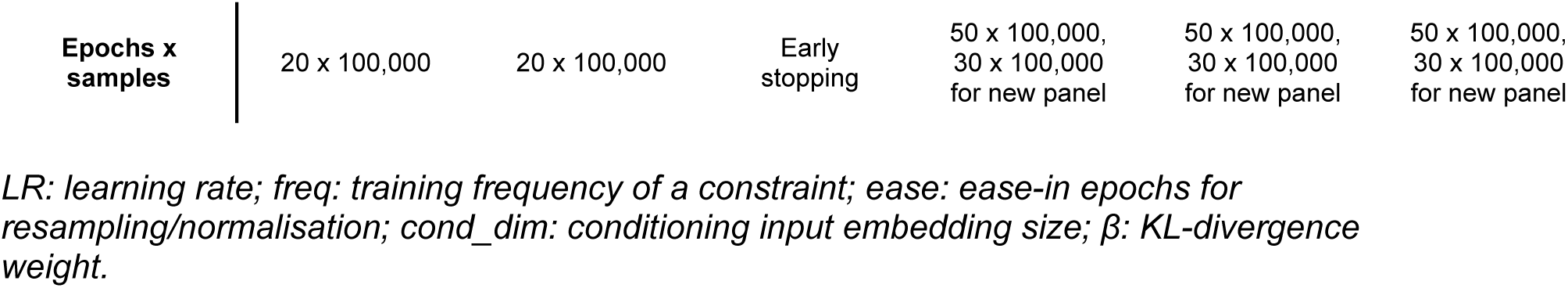
Summary of UVAE configurations for key experiments. This table details the specific components and key hyper-parameters of the UVAE framework used for the main analyses presented in the Results section.

The results demonstrate that a model incorporating all components: conditioning, latent normalisation, Subspace merging, MMD, and class-balanced resampling, achieved the best overall performance, with the highest batch mixing and lowest cluster mixing scores (Table 2, rank 1, full results in Table S1). Models that included the resampling component consistently outperformed their non-resampled counterparts, highlighting the importance of balancing class proportions during integration. The two most significant components for high performance were latent space normalisation for batch correction and Subspace merging for panel integration, which were present in all top-performing configurations. While combining all constraints yielded the best alignment, we observed a trade-off, as the model with the best imputation accuracy did not include the MMD merging constraint (Table 2, rank 5). These findings suggest that an integrated model with multiple, concurrent constraints results in the best performance, but trade-offs between alignment and generation accuracy must be carefully weighted. The weighting can be determined automatically by considering appropriate alignment metrics, chosen by the user for a specific problem.

**Table 2.** Best performing combinations of UVAE components on the synthetic dataset. Configurations are ranked by overall performance across three metrics: imputation MSE, batch mixing (iLISI), and cluster separation (cLISI). LISI scores are calculated in the latent space. Full results are available in Table S1, and full model configuration in Table 1.

| Conditioning<sup>a</sup> | Latent Norm. | MMD<sup>b</sup> | Subspace | Resampling<sup>c</sup> | Overall Rank | Imputed MSE (median/rank) | iLISI (median/rank) | cLISI (median/rank) |
| --- | --- | --- | --- | --- | --- | --- | --- | --- |
| ✓ | ✓ | ✓ | ✓ | ✓ | **1** | .29 / **3** | 6.37 / **1** | 1.27 / **1** |
|  | ✓ | ✓ | ✓ | ✓ | **2** | .29 / **2** | 6.26 / **2** | 1.28 / **2** |
|  | ✓ |  | ✓ | ✓ | **3** | .29 / **4** | 5.70 / **4** | 1.29 / **4** |
|  |  | ✓ | ✓ | ✓ | **4** | .32 / **7** | 5.74 / **3** | 1.29 / **5** |
| ✓ | ✓ |  | ✓ | ✓ | **5** | .28 / **1** | 4.69 / **17** | 1.28 / **3** |

<sup>a</sup>Conditioning applied to both encoder and decoder.
<sup>b</sup>MMD applied between panels only.
<sup>c</sup>Resampling using predicted cell type to balance latent normalisation and MMD.

We also found that specific architectural choices were important for preventing over-alignment. Conditioning both the encoder and decoder with the batch identifier, as opposed to only the decoder, reduced the risk of undesirable class mixing and loss of imputation accuracy, particularly when a linear embedding layer was used for the conditioning variable (Figure 3). This is a crucial consideration for real-world datasets with many batches, where such an embedding layer may be necessary to reduce the dimensionality of the conditioning vector.

**Figure 3.**
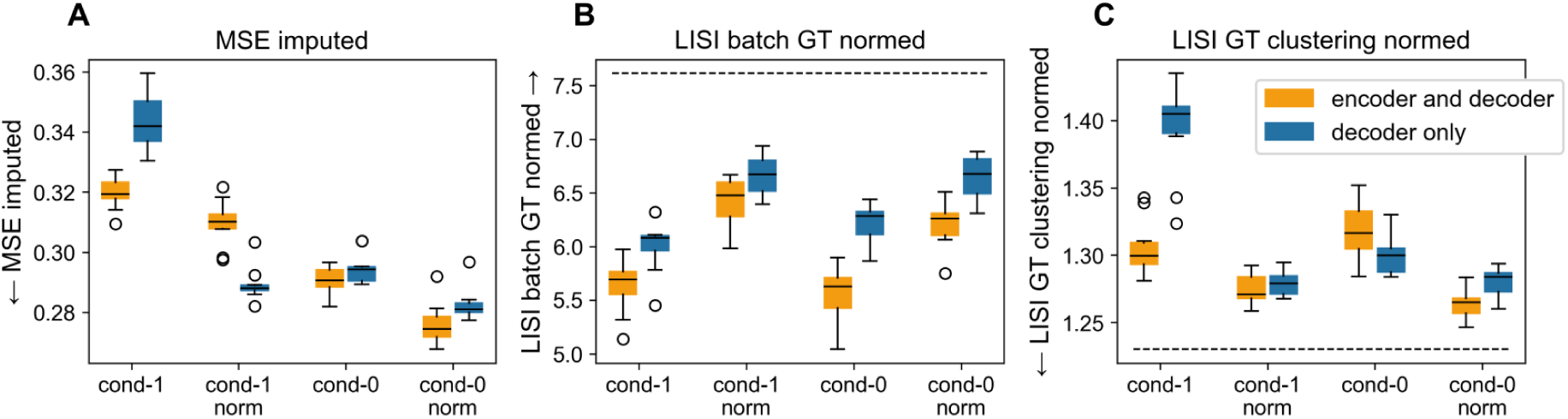
Encoder and decoder conditioning mitigates over-alignment. Comparison of model performance when conditioning both the encoder and decoder versus the decoder only. Results are shown for models with or without latent space normalisation (norm) and with either direct concatenation of the conditioning variable (cond-0) or a linear embedding (cond-1). (A) Imputed MSE. (B) Normalised batch LISI score (higher is better). (C) Normalised ground-truth cluster LISI score (lower is better). Dotted lines indicate scores for a ground-truth model trained on unaltered data.

### Batch clustering can be used to prevent over-alignment of unlabelled data

Our results show that resampling with class predictions obtained from expertly annotated data significantly improves performance (Table S1), however, sometimes annotations can be unavailable, or insufficiently detailed to allow for balancing of smaller class subpopulations. For unlabelled data, we developed an unsupervised strategy combining independent batch-level clustering for both resampling and alignment scoring.

For datasets lacking sufficient labels to train a reliable classifier, clustering is performed independently on each batch of data (with a user-chosen method of clustering, such as GMM). These batch-specific cluster assignments are then used as stand-ins for class labels. During resampling, an equal number of cells is drawn from each cluster within each batch. This approach does not assume that clusters are shared across batches and instead enforces a uniform local density within each batch’s latent representation, which helps to prevent over-mixing of disparate data regions during alignment. Importantly, our strategy does not require that the batch effects are small (in which case we could simply cluster all the data together to create shared class assignments).

For calculating unsupervised LISI alignment metric, we first showed that a correctly aligned model results in a cluster alignment score equal approximately to the number of integrated batches (since the batches overlap, one unique cluster assignment from each batch is expected in each neighbourhood), providing a valid optimisation target distinct from over-separation or random over-mixing (Figure S2A).

We then demonstrated the impact of unsupervised resampling combined with unsupervised LISI optimisation on model performance (Figure S2B-D). Models trained with resampling showed improved imputation accuracy, with lower MSE across all configurations (Figure S2B). A minor decrease in batch alignment was observed (Figure S2C), with a potential substantial increase in ground-truth class separation for models using conditioning (Figure S2D). Further aiming to enhance the stability of this unsupervised approach, we explored using multiple parallel clusterings for resampling. While this led to modest further improvements in imputation accuracy for some model configurations, the benefit was not consistently significant in our tests (Figure S3), suggesting that a single, well-chosen clustering is often sufficient to enhance integration accuracy.

We further investigated the performance gap between the supervised and unsupervised approaches. The supervised models require a classifier for resampling and LISI calculation, training of which may help to align the latent space by training the encoders. To better understand the value of unsupervised LISI scoring, we trained a classifier in both supervised and unsupervised cases but did not use its predictions for resampling in either case. In the supervised case, the classifier is used after training to calculate class-normalised LISI scores.

The results of this isolated test are shown in Table S2 and Figure S4. We observed that using the unsupervised LISI objective does not significantly lower either imputation or alignment performance compared to supervised cases, demonstrating its utility when detailed labels are unavailable.

### UVAE improves single-cell imputation accuracy in benchmark comparisons

We compared our method against cyCombine^6^, a state-of-the-art approach designed for the integration of flow cytometry data. We evaluated both distributional alignment and single-cell imputation accuracy. The detailed UVAE configurations used are shown in Table 1. For distributional alignment, we used the Dana-Farber Cancer Institute dataset (*DFCI*, consisting of two separate panel evaluations, and one evaluation of a batch shared across the two panels) and *van Gassen* dataset as per cyCombine publication, and we followed their evaluation methods by calculating Earth Mover’s Distance (EMD) for batch mixing and Mean Absolute Deviation (MAD) for preservation of biological variance. UVAE demonstrated stronger batch alignment on three of the four test scenarios, achieving higher EMD scores than cyCombine (Table 3). This came at the cost of higher MAD scores, indicating a greater reduction of variance, a characteristic trade-off for VAE models that regularize the latent space.

**Table 3.** Comparison of distributional alignment with cyCombine on public datasets. Batch alignment (EMD) and preservation of biological variance (MAD) were compared on the Dana-Farber Cancer Institute (DFCI) and van Gassen datasets. Higher EMD scores indicate better batch mixing, while lower MAD scores indicate less variance reduction. Bold values indicate the better-performing method for each metric and dataset.

| Dataset | EMD (cyCombine<sup>a</sup>) | EMD (UVAE) | MAD (cyCombine<sup>a</sup>) | MAD (UVAE) |
| --- | --- | --- | --- | --- |
| DFCI1 | 0.66 | **0.91** | **0.02** | 0.11 |
| DFCI2 | 0.66 | **0.94** | **0.02** | 0.18 |
| DFCIb3 | **0.86** | 0.74 | **0.01** | 0.07 |
| van Gassen | 0.84 | **0.90** | **0.02** | 0.20 |

<sup>a</sup>Scores from the original cyCombine<sup>6</sup> publication.

While distributional metrics assess batch-level alignment, they are insensitive to the accuracy of integration at the single-cell level. To address this, we designed an imputation task on the DFCI and van Gassen datasets. We computationally split each dataset into three panels and randomly held out five markers from each panel. We then trained models to impute these missing markers and measured the imputation performance. Since we expect the ground-truth values to contain batch effects, we measured the performance using Spearman’s rank correlation, which is robust to monotonic transformations such as shifts and scaling.

The results show that UVAE achieves substantially higher imputation accuracy than cyCombine across all tested datasets (Table 4). For the DFCI datasets, a UVAE model incorporating latent space normalisation, conditioning, and resampling performed best, with performance further improving when the correct target batch was specified during data generation. For the van Gassen dataset, a simpler UVAE model using only Subspace constraints for merging resulted in the highest accuracy.

**Table 4.** Comparison of single-cell imputation performance on public datasets. The mean Spearman correlation between imputed and original marker values is shown. Higher values indicate more accurate single-cell imputation.

| Model | Imputation target | DFCI1 (Spearman R) | DFCI2 (Spearman R) | van Gassen (Spearman R) |
| --- | --- | --- | --- | --- |
| **UVAE, sub. only** | non-spec. | 0.487 | 0.538 | **0.569** |
| **UVAE, sub. cond. norm. res.** | mean | 0.496 | 0.544 | 0.572 |
| **UVAE, sub. cond. norm. res.** | known batch | **0.517** | **0.562** | 0.543 |
| cyCombine, 4x4 SOM | non-spec. | 0.317 | 0.364 | 0.427 |
| cyCombine, 8x8 SOM | non-spec. | 0.319 | 0.366 | 0.437 |
| cyCombine, 12x12 SOM | non-spec. | 0.321 | 0.370 | 0.436 |
| cyCombine, 16x16 SOM | non-spec. | 0.321 | 0.370 | 0.292 |

This finding is consistent with the analysis on our synthetic dataset, where the ground-truth is known and imputation error can be measured directly with MSE. On that dataset, UVAE’s imputation error was approximately half that of cyCombine (Table 5), for both supervised and unsupervised UVAE variants. These results suggest that metrics like EMD and MAD may not fully capture the quality of data integration at the level of individual data points. UVAE’s generative approach, which learns cell-specific representations, provides a more accurate reconstruction of heterogeneous datasets at the single-cell level, which preserves the integrity of individual cellular profiles for downstream analyses.

**Table 5.**
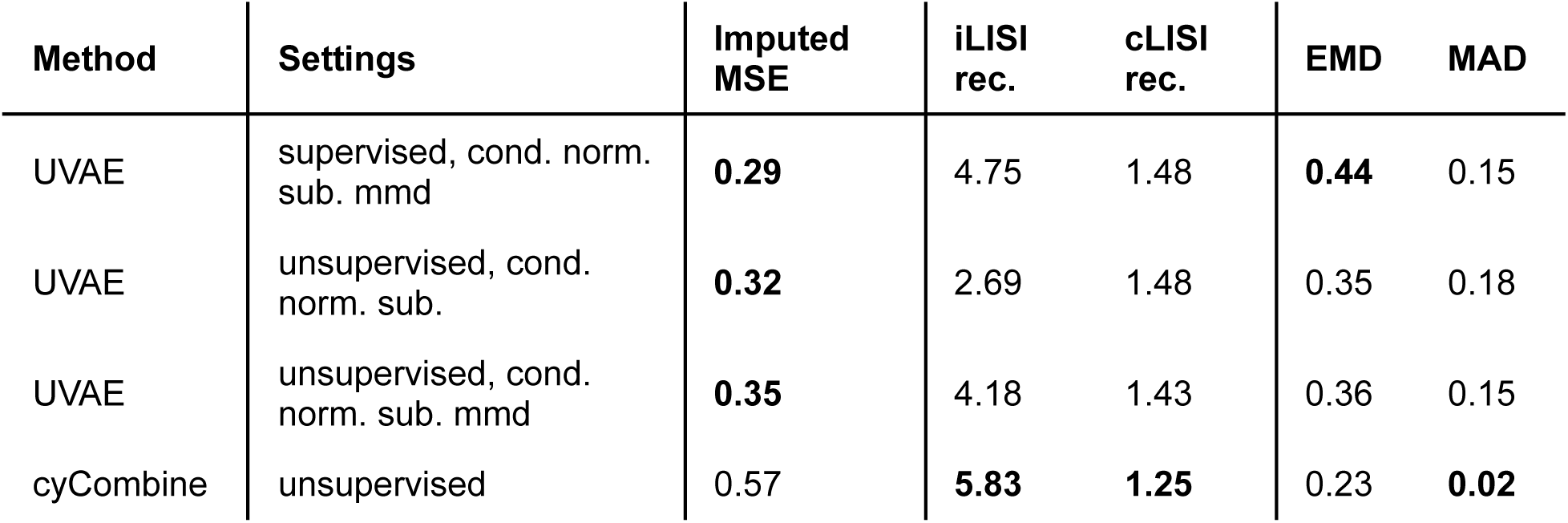
Comparison with cyCombine on synthetic data. Performance was measured using distributional metrics (iLISI, cLISI, EMD, MAD) and single-cell level accuracy (Imputed MSE). All metrics were computed on the reconstructed data space after integration and imputation.

| Method | Settings | Imputed MSE | iLISI rec. | cLISI rec. | EMD | MAD |
| --- | --- | --- | --- | --- | --- | --- |
| UVAE | supervised, cond. norm. sub. mmd | **0.29** | 4.75 | 1.48 | **0.44** | 0.15 |
| UVAE | unsupervised, cond. norm. sub. | **0.32** | 2.69 | 1.48 | 0.35 | 0.18 |
| UVAE | unsupervised, cond. norm. sub. mmd | **0.35** | 4.18 | 1.43 | 0.36 | 0.15 |
| cyCombine | unsupervised | 0.57 | **5.83** | **1.25** | 0.23 | **0.02** |

### Application to heterogeneous COVID-19 clinical data reveals severity-associated immune signatures

We next applied UVAE to integrate clinical flow cytometry data from the CIRCO consortium, collected from COVID-19 patients and healthy controls (Table 6). The dataset was highly heterogeneous, comprising 12 distinct marker sets for the whole blood lineage panel and 18 for the neutrophil-focused chemokine panel. We used UVAE to harmonise this data, correcting for batch effects and imputing a complete, uniform marker set for all cells. The integration strategy for both the lineage and chemokine panels, including the specific UVAE components used, is summarized in Table 1, with further details on hyper-parameter selection available in the STAR Methods.

**Table 6.**
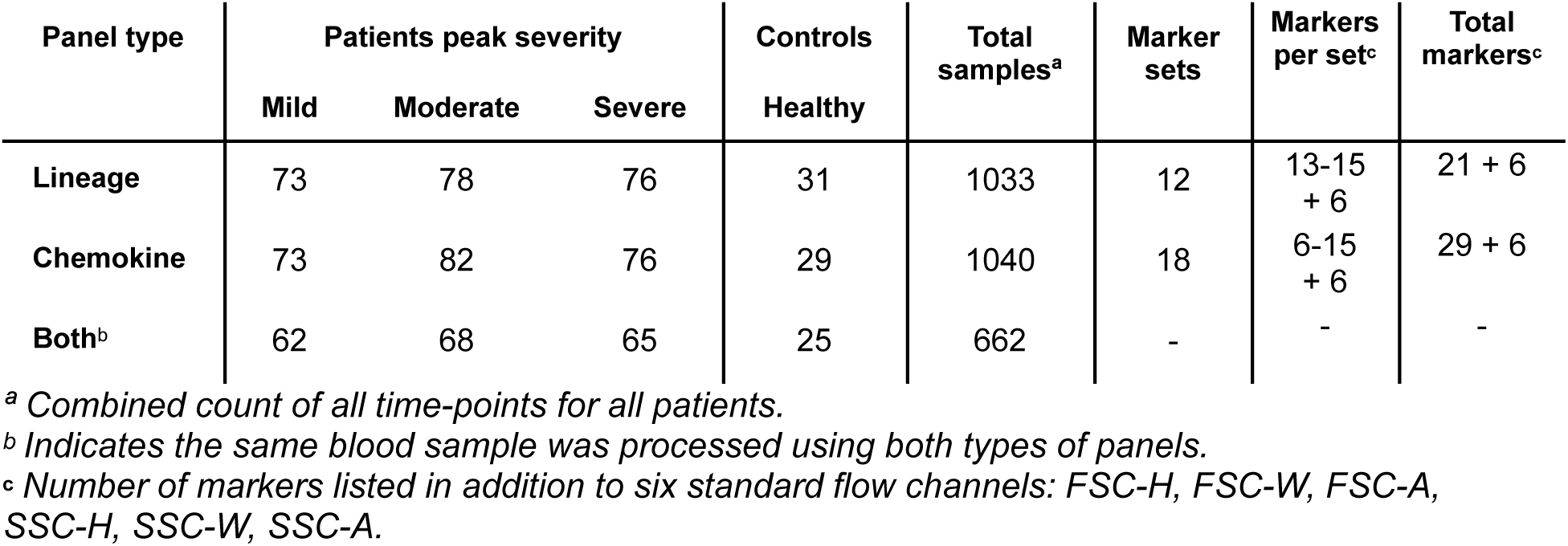
Patient and flow sample counts by panel type and peak clinical severity. The table shows the total number of patients and flow cytometry samples included in the study, categorized by the flow cytometry panel type used for analysis and the peak observed clinical severity.

For the whole-blood lineage panel type, we integrated 12 marker sets into a unified 50-dimensional latent space. UMAP visualisation of this space demonstrates successful alignment of data from different original panels, while preserving clear separation of manually annotated cell types (Figure 4A, 4B). We then regenerated a uniform marker set for all cells, correcting for batch effects. The resulting homogeneous data space is well-structured and suitable for downstream analysis (Figure 4C). A visualization of all imputed marker values is shown in Figure S5.

**Figure 4.**
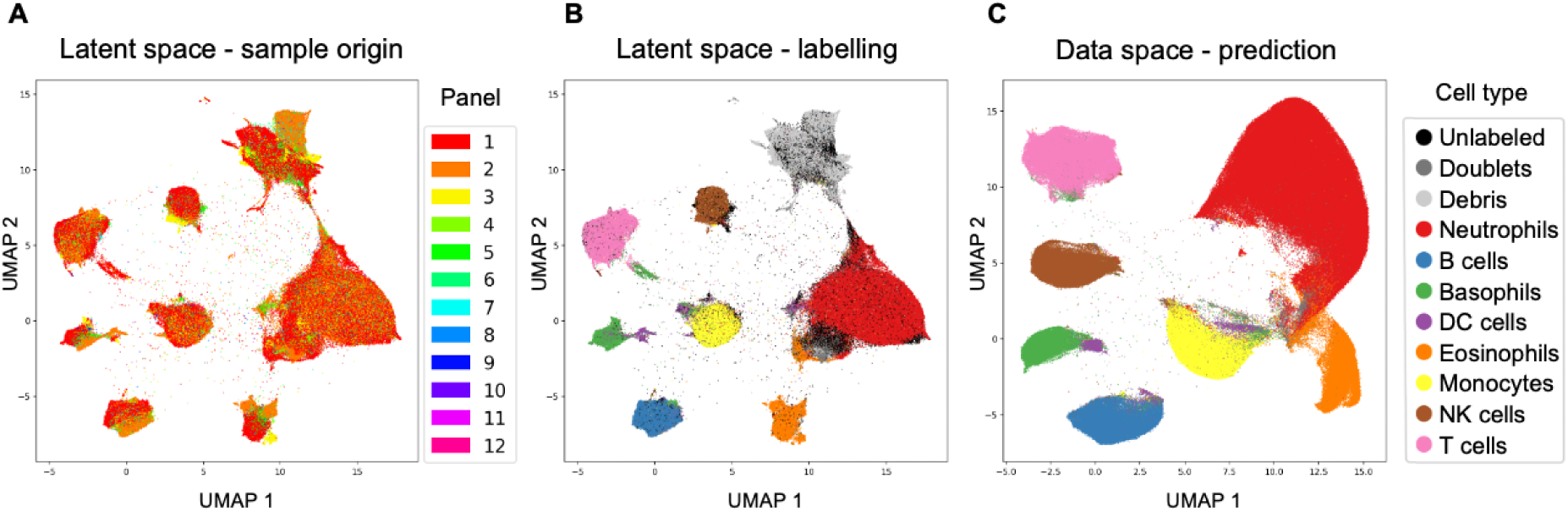
Integration of whole blood lineage panels from the COVID-19 dataset. (A) UMAP visualisation of the integrated latent space, coloured by the original marker set (*Panel* overlap), showing data alignment. (B) The same latent space coloured by manual cell-type annotation, demonstrating preservation of biological structure. (C) UMAP of the reconstructed data space after batch effect correction and imputation. Cell type, debris, and doublets annotations were predicted for all cells, and debris were excluded. Cells coloured by their predicted cell type.

We used the integrated data to compute cell-type proportions for all patient time-point samples, grouped by peak patient severity shown in Figure 5. Full data is shown in Table S3 for acute samples, and Table S4 for follow-up samples. In acute COVID-19 samples, we observed a significant increase in the proportion of neutrophils with escalating disease severity, alongside a corresponding decrease in T cells, DC cells, NK cells, and basophils, consistent with the known signatures of neutrophilia and lymphopenia in severe COVID-19^15,16^. We also found a significant decrease in eosinophils, which has been associated with poor prognosis^17^. When comparing these results to proportions derived from manual gating of the original data, the statistical significance of the associations with disease was substantially strengthened for nearly all cell types after UVAE integration (Table 7). For example, the p-value for the association between neutrophil proportions and disease status was five orders of magnitude smaller. In follow-up samples from recovered patients, most cell-type proportions returned to baseline, and we did not detect significant (q < 0.05 after Benjamini-Hochberg correction) differences in cell-type proportions across controls and patients grouped by severity.

**Figure 5.**
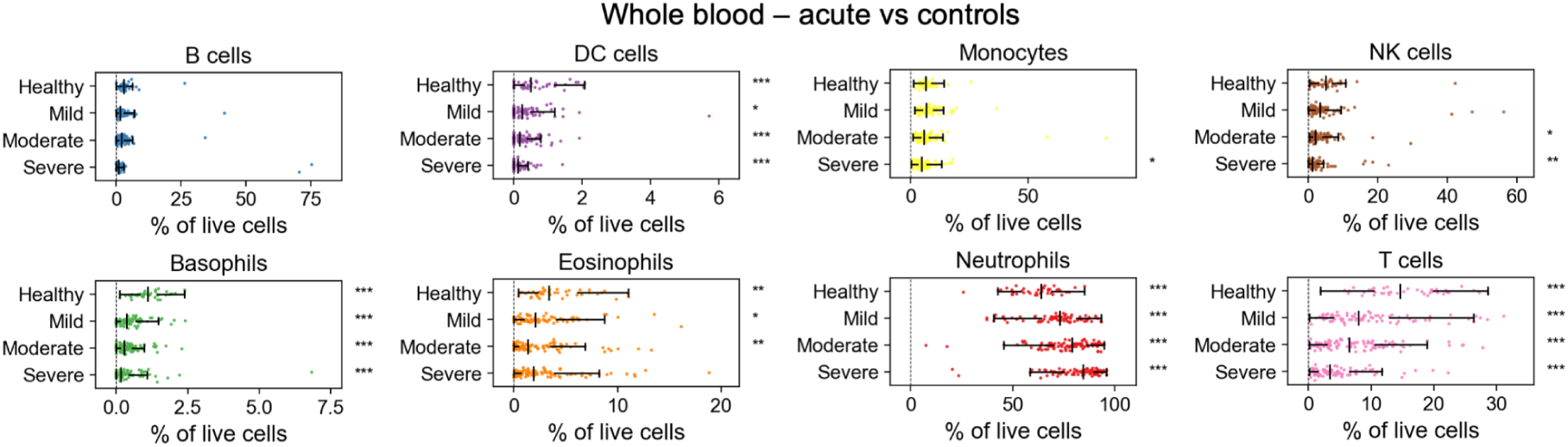
Cell-type proportions in whole blood correlate with COVID-19 severity. Proportions of major immune cell types are shown for healthy controls and patient samples, grouped by peak clinical severity. Significance indicators (*q<0.05, **q<0.01, ***q<0.001) shown to the right of each row were obtained from Welch’s t-test after Benjamini–Hochberg correction. Indicators on the Healthy row refer to the significance between healthy samples and all patient samples pooled. Mild/Moderate/Severe row indicators refer to each severity individually compared to healthy samples.

**Table 7.**
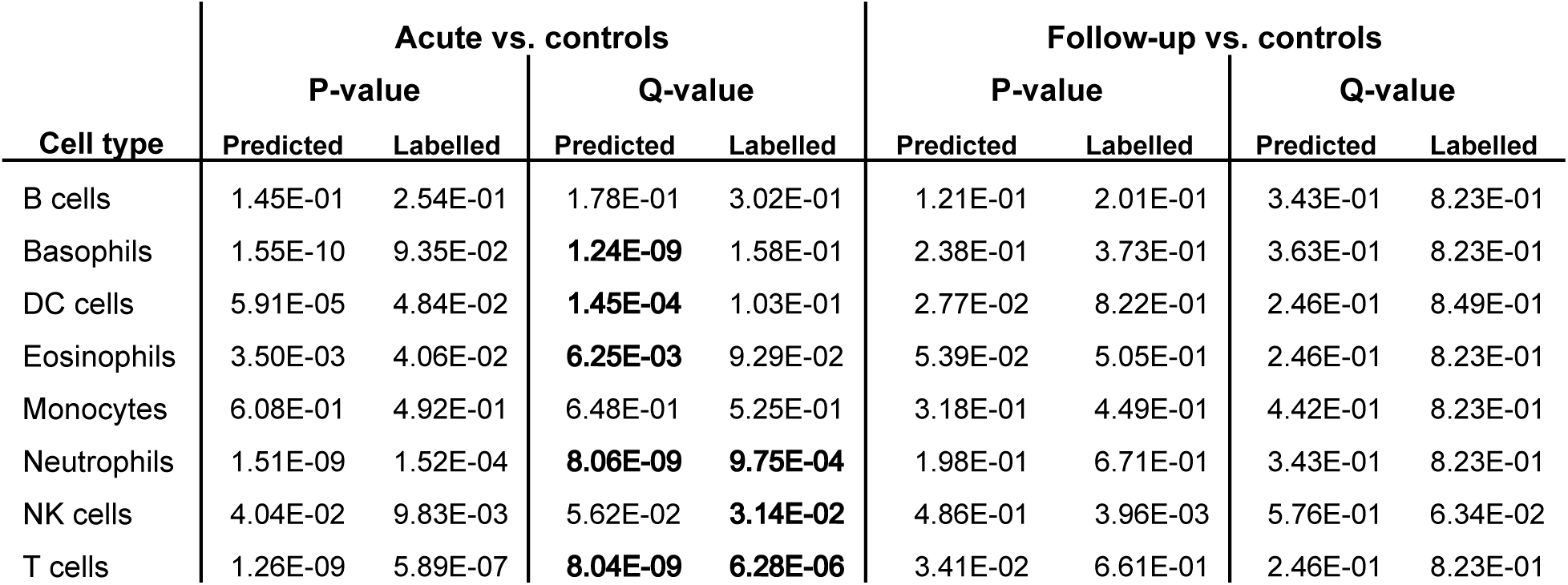
Comparison of statistical significance for cell-type proportions using UVAE predictions versus manual labels. The table shows raw p-values and Benjamini-Hochberg adjusted q-values from Welch’s t-tests comparing cell-type proportions between patient samples and healthy controls. Comparisons were performed separately for all acute samples and all follow-up samples, with p-values adjusted across the family of 8 cell types with 4 severity comparisons in each (see Table S3 and S4 for full data). Bold values indicate statistical significance (q < 0.05).

For the chemokine panel type, we integrated 18 marker sets into a common latent space and filtered the predicted non-doublet neutrophil subset (Figure S6 and S7). We then investigated the expression of activation markers on neutrophils (Figure S7). After imputation and batch correction, we found that the levels of several markers, including CCR2, CD49d, CX3CR1, and CD64, were significantly correlated with increasing disease severity in both acute and follow-up samples (Figure 6 and Table S5). CD86 was the most strongly associated marker in the acute phase. The homogeneity of the integrated data also enabled robust clustering of the neutrophil population. Using a GMM, we identified five distinct neutrophil clusters based on their marker profiles (Figure S6A). Figure S8 shows marker association with clusters, and Figure S9 shows cluster associations with disease severity. One cluster, characterised by low expression of multiple activation markers (including CD16, CD62L, and CD86), was significantly enriched in patients with severe disease. This finding points towards the presence of dysfunctional or immature neutrophil populations, a hallmark of severe COVID-19^17–19^.

**Figure 6.**
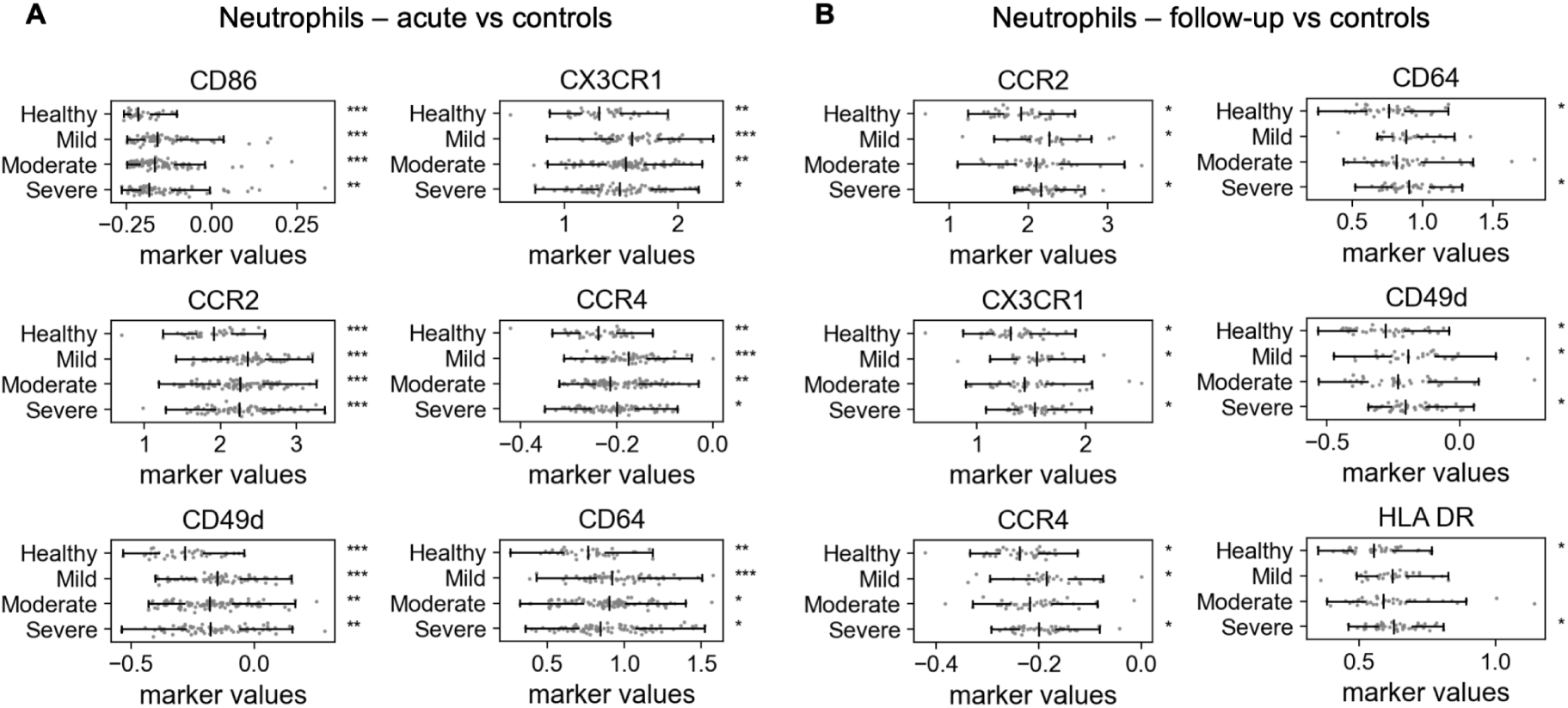
Neutrophil activation markers correlate with COVID-19 severity. Expression levels of key markers on neutrophils from the integrated chemokine panel data. Six most significant markers shown in each group. See Table S5 for a full list of markers. (A) Comparing acute samples to controls. (B) Comparing follow-up samples to controls. Significance indicators (*q<0.05, **q<0.01, ***q<0.001) shown to the right of each row were obtained from Welch’s t-test after Benjamini–Hochberg correction of all marker/severity comparisons within each group (acute or follow-up). Indicators on the Healthy row refer to the significance between healthy samples and all patient samples pooled. Mild/Moderate/Severe row indicators refer to each severity individually compared to healthy samples.

### Integrated longitudinal data improves prediction of peak COVID-19 severity

The unified dataset generated by UVAE provides a consistent foundation for building predictive models. We next sought to determine whether this integrated data could be used to predict a patient’s peak clinical severity from a longitudinal series of blood samples. We tested several model architectures using either cell-type proportions or raw marker expression values (*set-based* models) as input. For proportions, we used predicted cell-types of the whole-blood panel, and clustering of the neutrophil population of the chemokine panel. We investigated coarser GMM (5 clusters, see example in Figure S6A) and finer Leiden (varying number of clusters depending on the cross-validation fold, see example in Figure S6B) clustering assignment. We only used patient samples with both lineage and chemokine samples available, to evaluate the effects of including one or both types of panels. This resulted in a set of 26 healthy, 41 mild, 47 moderate, and 34 severe longitudinal samples.

To ensure a rigorous evaluation, we implemented a 4-fold cross-validation scheme at the patient level, where for each fold, a complete UVAE integration model was trained from scratch using only the data from the remaining three folds. Afterwards, the held-out data was projected onto the previously integrated space. In each case cluster assignments were defined on the training data and then transferred to the test data using a K-nearest neighbour classifier. This procedure prevents any information from the test set from influencing the data integration process. As a baseline, we calculated the MSE and F1 scores expected from a random classifier (randomly choosing a class with equal probability) and a stratified classifier (randomly choosing a class given known class probabilities).

The results demonstrate that set-based models operating directly on the integrated marker values consistently outperformed those using proportions (Table 8). Set-based models achieved both lower regression error (MSE) and higher classification accuracy (F1 score) across nearly all configurations. Increasing the number of cells sampled per time-point from 20 to 100 improved classification performance for the most complex models. The best overall classification score (avg. F1 = 0.466) was achieved by a set-based model with 100 cells per timepoint, using both whole-blood and chemokine panel inputs, and an RNN to pool temporal information. This highlights the complementary predictive power of the two panels and the value of modeling temporal dependencies. Notably, simpler linear models struggled to predict severe cases, underscoring the need for non-linear architectures to capture the complex immune signatures of disease.

**Table 8.**
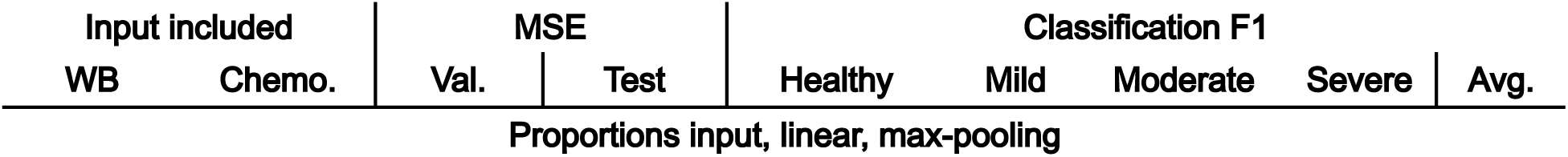
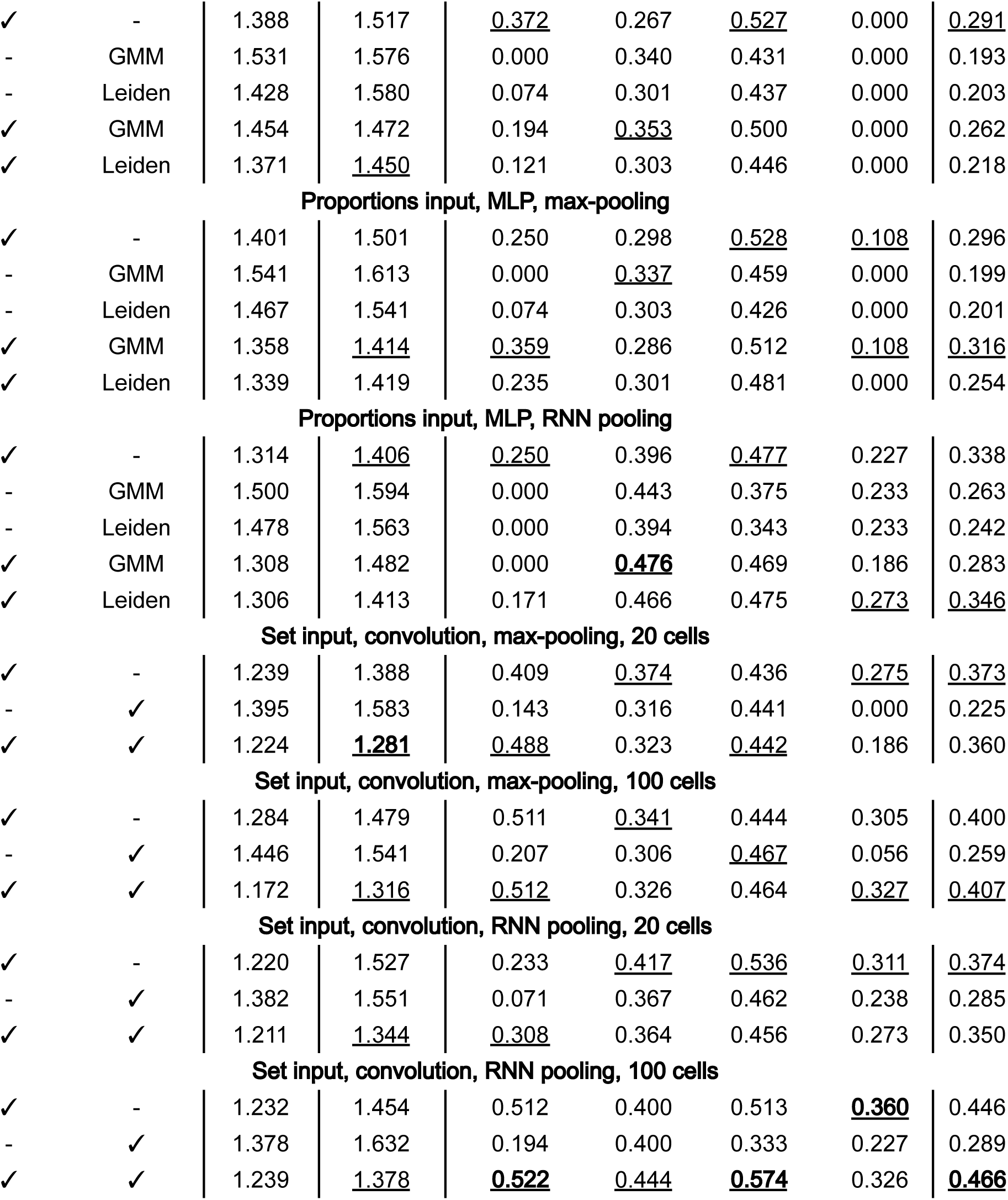
Cross-validation performance in regression of peak severity for longitudinal samples. MSE shows validation and test losses for ordinal regression. F1 scores are calculated by rounding regression output to assign a severity class. Underlined scores are best in each model category, bold scores are best overall. Expected scores from a uniform random classifier: MSE: 3.895, F1 Healthy/Mild/Moderate/Severe/Average: 0.206/0.263/0.280/0.239/0.247; stratified random classifier: MSE: 3.508, F1s: 0.176/0.277/0.318/0.230/0.250. WB – whole blood lineage input; Chemo. – neutrophil input from chemokine panel; MLP – multilayer perceptron; RNN – recurrent neural network.

For interpretability, we used the globally integrated dataset (trained on all data) to visualize patient trajectories and perform feature attribution. We used a max-pooling set model to predict the severity of individual time-points, which allows us to interpret the severity of each longitudinal sample with detail beyond the existing label of peak severity (Figure 7). Feature attribution analysis with a model trained only on chemokine data identified neutrophil markers CCR2, PDL1, CX3CR1, and CD64 as being among the most predictive features for determining severity, consistent with our earlier statistical analysis and the known immunopathology of COVID-19 (Figure S10). These results show that the homogeneous data generated by UVAE can be used to train powerful, interpretable models for tracking and predicting clinical outcomes from complex longitudinal data.

**Figure 7.**
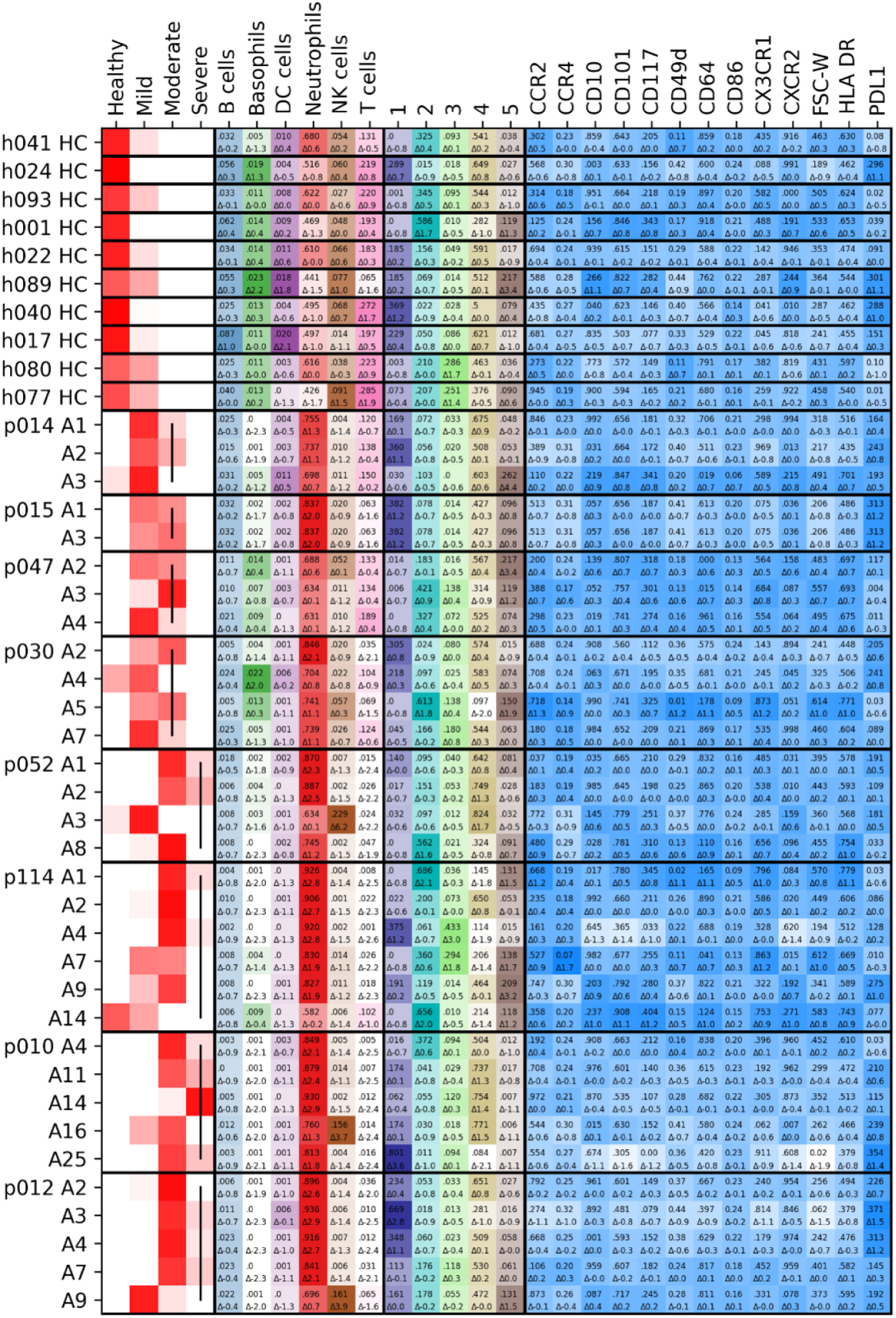
Visualisation of predicted disease trajectories for representative patients. Figure compares healthy controls (HC) with longitudinal samples from COVID-19 patients (p…). Columns show, from left to right: predicted severity for each time-point, whole-blood cell-type proportions, neutrophil cluster proportions, and key neutrophil marker values. Patient rows are ordered by time since hospital admission. The top number in each field is the value; the bottom number is the standard deviation from the control mean, also indicated by colour intensity.

## Discussion

The integration of heterogeneous clinical data presents significant challenges, including disparate feature sets, batch effects, and biological variability between samples. We have presented the Unbiasing Variational Autoencoder, a framework that addresses these issues through a modular, multi-VAE architecture. By applying UVAE to clinical flow cytometry data from COVID-19 patients, we demonstrated that the integration and imputation of a complete, harmonised dataset enhances the statistical power for identifying disease-associated biomarkers, enables robust clustering of cell subpopulations, and enables the training of downstream models for predicting clinical outcomes without resorting to removal of incomplete data.

Our benchmarks demonstrate that while methods like cyCombine can achieve strong alignment at the level of a distribution, UVAE provides superior imputation accuracy, signifying that more information is captured by the model at the single-cell level. Metrics like EMD and MAD are not sensitive to permutations and do not capture the fidelity of reconstruction for individual cells. UVAE’s generative nature, which learns a specific latent representation for each cell, results in more accurate imputations, as shown by both higher Spearman correlations on public datasets and lower MSE on our synthetic ground-truth data. We observed that the strong alignment of VAE-based models can come at the cost of reduced variance in the reconstructed data. This appears to be a fundamental trade-off of VAEs that regularize the latent space; using a non-variational autoencoder in our framework (equivalent to β=0) preserves variance but results in poor alignment, whereas a standard VAE (β=1) effectively aligns data at the cost of this variance reduction.

The flexibility of the UVAE framework requires user consideration of several design choices. We consistently used a latent dimensionality of 50, a value greater than the number of markers in any single panel (27 for lineage, 35 for chemokine), to ensure the latent space was sufficiently expressive. This avoids forcing information compression through a bottleneck, instead allowing regularisation to be driven by the VAE’s Gaussian prior and the alignment constraints. The choice of MMD for density matching was based on its demonstrated effectiveness in similar VAE-based batch correction tasks^10,11^ and its suitability for Gaussian latent spaces, though the framework is flexible and can accommodate other distance metrics. Similarly, we chose GMM for unsupervised resampling for its speed and predictability; users should apply domain knowledge and can select other methods, such as Leiden clustering or geometric sketching^20^, that are best suited to reveal meaningful biological structure in their data. Hyper-parameter optimisation may also require iterative refinement, as users may need several attempts to determine the meaningful range for alignment scores like LISI, thereby controlling the trade-off between alignment and reconstruction quality.

For clinical applications, the reliance on automatically merged and imputed data requires careful consideration. Unsupervised clustering may obscure rare but biologically important cell populations, and over-regularization could remove subtle sources of variation. Therefore, automated outputs should be inspected to ensure they align with expert knowledge. While the model is capable of working with no, full, or partial labelling, our results confirm that expert manual annotation is always beneficial, as demonstrated by the superior performance of supervised models over unsupervised ones.

A further consideration is the handling of model uncertainty. Our approach, like many multi-stage analysis pipelines, uses the generative model to produce single point estimates for imputed values, and this imputation uncertainty is not propagated to downstream statistical tests or predictive models. This could lead to an overestimation of statistical significance. While VAEs, as generative models, could in principle quantify this uncertainty on a per-cell basis, its practical application is non-trivial. Standard downstream tools are not designed to incorporate such error estimates. More fundamentally, the VAE provides independent uncertainty for each cell, which does not capture the correlated errors needed to correctly estimate the uncertainty of aggregate statistics like cell-type proportions or mean marker levels. Developing methods to model and propagate this structured uncertainty is therefore an important area for future research.

Training of the UVAE models is computationally efficient, on our dataset requiring approximately 20 minutes for the initial three-panel model on a single CPU. The cost scales mainly with the number of samples included in training, and number of constraints in the model (see Table S7 for training and inference times). Additionally, hyper-parameter optimisation can significantly extend the time of model creation. As the number of integrated panels grows more autoencoders are needed, however, it becomes more likely that new data can be projected using an existing, pre-trained autoencoder, which can also be fine-tuned to accommodate new batches. Further avenues for optimisation remain; for instance, the hyper-parameters of the Mango optimizer itself were not explored. Through experience, we identified key hyper-parameters controlling the training dynamics, the *unsupervised* and *merge* learning rates, and the *pull* and *frequency* of individual constraints, and we advise users to focus on these for efficient optimisation.

In conclusion, this work introduces a significant methodological advance for the analysis of clinical flow cytometry datasets. UVAE’s principles of constrained latent space alignment and class-balanced integration extend to the analysis of any heterogeneous or unpaired data. Our Python framework is applicable to all real-valued data and easily extensible, providing a powerful tool for the integration of complex biomedical datasets.

## Resource Availability

● **Lead Contact:** Further information and requests for resources should be directed to and will be fulfilled by the Lead Contact, Magnus Rattray.
● **Materials Availability:** This study did not generate new unique reagents.
● Data and Code Availability:
  ○ The UVAE framework is available on GitHub: https://github.com/mikephn/UVAE.
  ○ The code used for training, benchmarking, and generating the results in this paper is available on GitHub: https://github.com/mikephn/UVAE-COVID19-codebase.
  ○ The CIRCO COVID-19 flow cytometry data and the synthetic benchmark data generated for this study are available at the following Zenodo repository: http://doi.org/10.5281/zenodo.13854783. All other publicly available datasets used for benchmarking are cited with their respective accession codes in the Key Resources Table.

## Funding

This work was supported by The Kennedy Trust for Rheumatology Research that provided a Rapid Response Award for costs associated with the laboratory analysis of the immune response in patients with COVID-19 to J.R.G., The Wellcome Trust (T.H., 202865/Z/16/Z; 106898/A/15/Z), UKRI-MRC (T.H., J.G, M.R. MR/V028448/1). The Oxford and Manchester NIHR BRC provided support for study design and sample collection.

## Author Contributions

Conceptualization: MP, JG, MR; data curation: MP, VK, TW, LL, JG; formal analysis: MP; funding acquisition: TH, JG, MR; investigation: MP; methodology: MP, MR; project administration: JG, MR; resources: JG; software: MP; supervision: JG, MR; validation: MP; visualization: MP; writing – original draft: MP, MR; writing – review & editing: all authors.

## Declaration of Interests

The authors declare no competing interests.

## Declaration of Generative AI

During the preparation of this work, the authors used Google Gemini 2.5 Pro to revise the text and Anthropic Claude 3 to document the software. The authors reviewed and edited the content and take full responsibility for the content of the publication.

## Supplemental Information Index

**Supplemental Information**

**Document S1. Algorithms S1-S8, Figures S1-S10, Tables S1-S7.**

## STAR Methods

### EXPERIMENTAL MODEL AND SUBJECT DETAILS (COVID-19 Patient Cohort)

Our clinical data consists of flow cytometry samples obtained between March 2020 and May 2021 from patients with COVID-19 admitted to four hospitals in the Greater Manchester, UK, region, as previously described^28^. All participants or their legal representatives provided written informed consent.

Multiple blood samples (between 1 and 6) were collected from patients throughout their hospitalisation and were labelled with the peak observed clinical severity (Mild, Moderate, or Severe) during their admission. Follow-up samples were collected from a subset of patients after recovery. Additionally, control samples were collected from healthy volunteers. Details of the patient cohort, including the number of patients and samples grouped by panel type and peak severity, are provided in Table 6 in the main text.

### METHOD DETAILS

#### UVAE Framework: General Overview

The Unbiasing Variational Autoencoder (UVAE) is a framework for automated training and optimisation of semi-supervised, VAE-based architectures. It is implemented in Python using the Keras library for neural network training. A UVAE model is composed of multiple, interconnected network components designed to match user-specified data inputs and alignment constraints.

The modular design allows for the specification of multiple, partially overlapping data and labelling series, which are used to define merging and normalisation constraints within a single generative model. The trained model can be saved and loaded, enabling persistence for subsequent analysis or the integration of new data onto a pre-existing model. The full training process is outlined in Algorithm S1.

The fundamental component of the framework is a β-VAE^29^ trained for each input data series that has a distinct feature set (e.g., a unique set of flow cytometry markers). All VAEs in a model share the same latent space dimensionality. The core architecture of each VAE consists of an encoder, which maps input data to a latent distribution (mean and log-variance), and a decoder, which reconstructs the data from a sample of this distribution. The model is trained to minimise the evidence lower bound (ELBO), which comprises a reconstruction loss term (mean squared error) and a regularisation term (KL-divergence between the latent distribution and a standard Gaussian prior), scaled by a hyper-parameter β.

Supervised annotations, such as cell-type labels, can be defined over arbitrary subsets of the data. These are used to train classification or regression functions that operate on the shared latent space (Algorithm S2). The predictions from these functions can be used to guide the integration process through class-balanced resampling. In addition to supervised prediction targets, annotations can be specified for merging or normalisation constraints, such as batch effect correction.

During training, mini-batches are defined for each trainable component (autoencoders, classifiers, merging constraints). These are combined and trained in a random order within each epoch. Gradient clipping is applied to enhance training stability. The total loss is the average of all mini-batch losses, including the ELBO, classification, regression, and merging losses. Three parallel Adam optimisers are used: one for unsupervised autoencoder training, one for supervised classifier/regressor training, and one for merging constraints. The relative learning rates of these optimisers can be tuned to balance the model’s performance across reconstruction, prediction, and data alignment objectives.

#### Batch Effect Correction Mechanisms

UVAE defines two methods of batch effect correction that can be applied concurrently within each VAE to separate the influence of nuisance covariates from the shared latent space. These mechanisms operate in addition to any MMD-based constraints that may also be used to minimise batch differences.

**Conditional Variational Autoencoding:** The first mechanism is conditioning, in the style of a Conditional VAE (CVAE^13^). For each cell, a categorical variable specifying the nuisance covariate (e.g., batch ID) is provided. This variable can be projected to a fixed dimensionality through a trainable linear embedding layer before being concatenated to both the encoder’s input data vector and the decoder’s input latent vector. The conditioning of both the encoder and decoder is recommended to mitigate over-alignment. This conditioning encourages the model to learn batch-specific encoding and decoding functions, thereby preventing the shared latent space from redundantly capturing this information. The nuisance covariate information is not passed to subsequent constraints that operate on the latent space, ensuring that downstream classifiers and merging functions receive a batch-agnostic representation.

**Latent Space Arithmetic:** The second mechanism is latent space normalisation, which directly offsets sample embeddings in the latent space. At the end of each training epoch, a mean latent vector is calculated for each batch using a class-balanced set of samples to ensure the representation is not biased by differing cell-type proportions. A common reference vector is then determined, which can either be the mean of a specific target batch or the average of all batch means. The offset required to move each batch’s mean to this common reference is then calculated. During subsequent training and inference, this pre-calculated offset vector is subtracted from the latent embedding of each cell originating from that batch. This process is introduced gradually during training via an *ease-in* hyper-parameter to maintain stability. The latent normalisation procedure is detailed in Algorithm S3. If multiple normalisation constraints are specified, their offsets are calculated and applied sequentially.

#### Integration of Disparate Panels

UVAE integrates data from disparate panels by aligning the latent space embeddings of their respective VAEs. This alignment is achieved through two complementary merging constraints that can be used simultaneously: feature matching through a Subspace constraint and density matching via MMD loss. The contribution of each constraint to the total model loss is controlled by a tuneable *pull* hyper-parameter.

**Feature Matching with a Subspace Constraint:** For data series that share a subset of input features (markers), a Subspace constraint provides a direct method for alignment. This constraint introduces an additional VAE that is trained only on the common features shared across all specified data series. During training, a data sample is encoded through both its panel-specific VAE and the shared Subspace VAE. Both encoders can perform their own batch effect correction, with the Subspace VAE correcting over a union of all batches from the panels it connects. The alignment is enforced by minimising the mean squared error (MSE) between the unbiased latent embedding from the panel-specific encoder and the unbiased latent embedding from the Subspace encoder for each sample. Since this distance is calculated on paired embeddings from the same cell, this process is robust to class imbalance between panels but can be sensitive to batch effects if not properly corrected. The Subspace merging process is detailed in Algorithm S4.

**Density Matching with MMD Loss:** To enable alignment without relying on shared features, or to further enforce alignment between panels, UVAE uses a density-matching approach. This is implemented by minimising the Maximum Mean Discrepancy (MMD) loss between the distributions of unbiased latent embeddings from different data series. MMD is calculated using a multi-scale Gaussian radial basis function (RBF) kernel, as described in trVAE^11^. For each training mini-batch, random samples are drawn from two data series (selected randomly if more than two are being merged). To prevent distortion from biological differences between the series, the samples used for MMD calculation are class-balanced using the resampling mechanism. The MMD loss is then back-propagated to the encoders of the involved VAEs, training them to produce more similar latent distributions. While primarily used for panel integration, MMD loss can also be applied as an additional batch effect correction mechanism within a single panel. The MMD merging process is detailed in Algorithm S5.

#### Class-Balanced Resampling

A key challenge in integrating heterogeneous datasets is that biological differences, such as varying cell-type proportions between batches or panels, can be misinterpreted as technical variation, leading to over-correction and distorted data alignment. UVAE addresses this through a class-balanced resampling mechanism that is applied to constraints sensitive to class imbalance, namely latent space normalisation and MMD-based merging. This component ensures that the distributions being compared are compositionally equivalent. The amount of resampled data is increased gradually during training via an *ease-in* hyper-parameter, which linearly increases the proportion over a specified number of epochs. To improve stability, multiple independent class assignments with different numbers of components can be applied in parallel, with each contributing to the resampling of a random portion of the data.

**Supervised Resampling:** In a semi-supervised setting where partial class labels (e.g., manually gated cell types) are available, a classifier is trained on the shared latent space to predict the class of every cell in the dataset. At the end of each epoch, these predictions are used to guide resampling. For a given constraint, the indices of the data are re-sampled to match a target class distribution. Typically, this target is the average proportion of each class across all batches or panels being integrated. This ensures that when batch means or MMD are calculated, each group is represented by a sample with the same underlying class composition, isolating technical variation from biological variation. The detailed procedure is outlined in Algorithm S6.

**Unsupervised Resampling with Batch-Level Clustering:** For datasets lacking sufficient labels to train a reliable classifier, an unsupervised resampling approach can be used. In this mode, clustering is performed independently on each batch of data (e.g., using a Gaussian Mixture Model [GMM] with a fixed number of components). These batch-specific cluster assignments are then used as stand-ins for class labels. During resampling, an equal number of cells is drawn from each cluster within each batch. This approach does not assume that clusters are shared across batches and instead enforces a uniform local density within each batch’s latent representation, which helps to prevent over-mixing of disparate data regions during alignment. The detailed procedure is outlined in Algorithm S7.

#### Generation of Homogenized Data

After training, the integrated UVAE model can be used as a generative tool to create a single, homogeneous dataset. This process of cross-panel translation imputes all missing marker values and corrects for batch effects across all cells, resulting in a unified data matrix that is suitable for direct comparison, clustering, and downstream statistical analysis.

The generation process leverages the aligned shared latent space as a common frame of reference. To generate a complete marker set for a given cell, the cell’s original data is first passed through its panel-specific encoder to obtain a latent representation. If batch correction mechanisms were used during training, they are first applied to translate the cell from its source batch to an unbiased latent space, and then in reverse to generate data in the style of a user-defined target batch. Specifically, if latent space arithmetic was used, the source batch’s latent offset is subtracted from the embedding, and the target batch’s offset is added. Similarly, if the VAEs were conditioned, the source batch ID is provided to the encoder, and the target batch ID is provided to the decoder.

This batch-translated latent vector is then passed through the decoder of a different VAE that was trained on a panel containing the desired markers. This decoder reconstructs the full set of markers for its panel, thereby imputing the missing values for the input cell. To enhance the stability and homogeneity of the final dataset, all cells are typically regenerated using a fixed set of decoders, usually those corresponding to the largest or most comprehensive panels. If a desired marker is present in multiple target decoders, the final imputed value is the arithmetic mean of the values generated by each. The full imputation process is detailed in Algorithm S8. This procedure transforms the original collection of heterogeneous, unpaired data streams into a single, coherent, and interpretable dataset.

#### Model Selection and Hyperparameter Optimisation

The performance of the UVAE framework depends on the selection of appropriate hyper-parameters, which control model architecture, learning rates, and the relative contributions of different loss components. Default hyper-parameters are shown in Table S6. For optimal performance on any given task, we conduct an automated hyper-parameter search using the Mango^25^ Bayesian optimizer to identify optimal settings that balance reconstruction accuracy, supervised prediction performance, and any custom data alignment metrics provided.

The optimisation objective function is a weighted sum of the model’s validation loss and one or more explicit alignment metrics. The validation loss itself is a composite of the evidence lower bound (ELBO), classification/regression losses, and merging losses, calculated on a held-out portion of the data.

**Alignment Metrics for Optimisation:** To directly measure the quality of data integration, we incorporate alignment scores into the optimisation objective. The primary metrics used are based on the Local Inverse Simpson’s Index (LISI)^24^, which quantifies the degree of mixing of labels within the local neighbourhood of each cell. We typically compute two LISI scores: an integration LISI (iLISI) to measure batch mixing (where higher scores are better) and a cell-type LISI (cLISI) to measure the preservation of biological separation (where lower scores are better). Alternatively, for direct comparison with other methods like cyCombine, we can use a combination of Earth Mover’s Distance (EMD) and Mean Absolute Deviation (MAD) as the alignment metrics.

**Normalised LISI Scoring:** LISI scores are sensitive to class imbalance between batches. To establish an empirical range for LISI scores and understand their behaviour on imbalanced data, we first trained worst-case models on our synthetic dataset. The maximum-separation model is trained on disjoint data with batch effects, without applying any form of correction or merging (Figure S1A). Subsequently, the maximum mixing model is obtained from this worst-case embedding by randomly permuting the locations for all samples (Figure S1B). We compared these to a ground-truth model trained on perfectly aligned data without batch effects (Figure S1C). Standard LISI scores, when applied to these models, proved to be misleading on datasets with imbalanced class compositions across batches. As shown in Figure S1D, a naively over-mixed (shuffled) embedding can receive an iLISI score higher than the ground-truth model, because in the ground-truth the batches are not ideally uniformly distributed through the latent space.

To address this, we define a normalised LISI score for supervised scenarios. Before calculating LISI, we down-sample the cells in each batch to match the lowest count of each class present across all batches (Figure S1E illustrates ideal correction with known ground-truth classes, and Figure S1F a correction with imperfect labelling). This ensures that the score reflects the mixing of compositionally identical populations, providing a more accurate and reliable measure of batch integration that is not confounded by biological heterogeneity. We adopt this normalization for all supervised model selection.

**Unsupervised LISI Scoring:** In unsupervised scenarios we do not have access to class labelling or other assignments in which identities are shared across batches. Jointly clustering data with significant batch effects is unreliable, as the clusters tend to separate by batch rather than biology. We therefore define an unsupervised LISI score. In this approach, we first cluster each batch independently. The resulting batch-specific cluster IDs are then used as the labels for calculating a cLISI-like score to penalize over-mixing. In this context, a lower score is not necessarily better. With perfectly overlapping batches, the ideal score should be close to the number of integrated batches (provided high enough perplexity is used for LISI calculation), with one unique cluster coming from each batch.

We empirically validated this concept in Figure S2A. The ground-truth (GT) model’s unsupervised LISI score approaches the number of batches (nine) at higher perplexity values, clearly distinguishing it from a maximally separated model (score close to 1) and a randomly mixed model. This establishes an optimisation target for preserving biological separation in the absence of labels, guiding the model to align batches without erasing their unique internal structures.

**Hyper-parameter Search Space:** The Bayesian optimizer searches over a defined space of hyper-parameters, which can include VAE architecture (latent dimensionality, number of hidden layers, neuron counts), activation function slope, dropout rate, learning rates for the different optimizers, the *pull* strength of merging constraints, the β scaling factor for the KL-divergence, and the *ease-in* epochs for resampling and normalisation. Note that pull and frequency are optimized independently for every constraint. A full list of tuneable hyper-parameters and their default optimised ranges is provided in Table S6.

#### Application-Specific Implementations

This section provides the specific implementation details for the three main analyses presented in the Results: the validation of UVAE components using synthetic data, the benchmark comparison against cyCombine, and the application to the clinical COVID-19 dataset, including the downstream severity regression modeling. For each application, we describe the data pre-processing, the specific UVAE model configuration, and the evaluation procedure. A summary of the key model configurations for each experiment is provided in Table 1.

#### Synthetic Data Generation and Validation

To create a benchmark dataset with a known ground truth for evaluating alignment and imputation, we generated a synthetic dataset derived from a single, well-annotated flow cytometry sample. This approach ensures that any observed misalignment is purely a result of our computational manipulations, not underlying biological differences.

**Data Generation:** A single flow cytometry sample from a healthy control was selected from the CIRCO dataset. After removing debris and unlabelled events, a random subset of 100,000 cells comprising eight major immune cell types was taken and the marker values were standardised (mean=0, std=1). To create a ground-truth biologically meaningful assignment, more granular than the manual annotations, this data was clustered into 20 groups using a GMM. This cluster annotation was then used as the basis for creating artificial batch and panel effects. The dataset was split into three panels, and each panel was further split into three batches. The split was performed unevenly by assigning a disproportionate number of cells from each GMM cluster to each batch and panel, simulating biological heterogeneity. To create incompatible feature sets, two unique marker channels were removed from each of the three panels (CD45/CD11c from panel 1, CD16/CD11b from panel 2, CD10/CD14 from panel 3). Finally, the data within each of the nine resulting batches was independently re-standardised, introducing misalignments due to the varying cluster proportions.

**Validation Procedure:** The synthetic data was used to test various UVAE model configurations (see Table 1). Each model was trained to align the nine batches and three panels. Performance was assessed using three metrics: 1) imputation error, calculated as the MSE between the imputed marker values and the original ground-truth values; 2) normalised batch iLISI, to measure batch mixing; and 3) normalised ground-truth cluster cLISI, to measure preservation of the original GMM cluster structure. The imputation was performed for each panel by encoding its values using panel encoder, and decoding them using the remaining two decoders, while setting an arbitrary but constant target batch. The average of the two outputs was then standardised, and MSE loss calculated between the result and the original removed values. A ground-truth reference VAE model was trained on the unaltered, pre-split data to establish an ideal performance baseline. Hyper-parameter optimisation for each configuration was performed over 30 iterations to find settings that maximized a weighted combination of validation accuracy and the LISI alignment scores.

The ranges of LISI scores were defined by training two worst-case models (in the same way as described in *Normalised LISI Scoring* and Figure S1) to identify the top and bottom of expected score range. During optimisation, the actual score was placed in this range to be normalised between 0 and 1. The final reported metrics are the median scores from 10 models trained with the best-found hyper-parameters.

#### Benchmarking against cyCombine

We compared the performance of UVAE to cyCombine^6^ using two publicly available datasets from the original cyCombine publication and our own synthetic dataset. The evaluation was designed to assess both batch-level distributional alignment and single-cell level imputation accuracy.

**Dataset Preparation:** The Dana-Farber Cancer Institute (DFCI) dataset (FlowRepository ID: FR-FCM-Z52G), consisting of CyTOF data from healthy donors and patients with Chronic Lymphocytic Leukemia (CLL), was downloaded and prepared using the functions provided in the cyCombine package. This dataset includes two main panels with 15 shared markers. The van Gassen dataset (FlowRepository ID: FR-FCM-Z247) consists of 40 samples across 10 batches with 2 stimulation conditions and 37 measured markers.

**Distributional Alignment Benchmark:** We replicated the alignment benchmarks from the original cyCombine publication. For the DFCI data, this included batch correction within each of the two panels, and between the third batch from each panel. For the van Gassen data, this involved correcting across all 10 batches. For each scenario, a UVAE model was configured with VAE conditioning on batch and disease status, latent space normalisation of the batch, and MMD applied between batches. Resampling was performed using an unsupervised approach based on GMM clustering of the combined data. Hyper-parameters were optimised over 50 iterations to minimise a weighted combination of model validation loss, negative EMD score, and MAD score. The final EMD and MAD scores were calculated on the regenerated data using the cyCombine implementation. See Table 1 for configuration details.

**Imputation on Public Data:** To assess single-cell accuracy, we designed an imputation task for the DFCI1, DFCI2, and van Gassen datasets. Each dataset was randomly split into three panels at the sample level, ensuring no sample was spread across panels. From each panel, five markers were randomly selected and held out.

● **UVAE Application:** We tested several UVAE configurations (see Table 1). The best performing models for DFCI1 and DFCI2 included conditioning (of batch), latent normalisation (of batch), resampling (using 6 GMM clusters), and three Subspace constraints (one for each pair of panels). For the van Gassen dataset, a simpler model using only the three Subspace constraints performed best. Models were trained with default hyper-parameters, using 20% of the data for early stopping. For models with conditioning, we tested data generation using two target batch strategies: specifying the known correct batch for each cell, or averaging reconstructions across all shared batches to simulate an unknown target.
● **cyCombine Application:** To impute the missing channels for a given panel (e.g., panel 1), we first performed batch correction on it. We then concatenated the other two panels (panels 2 and 3) using their shared channels, applied batch correction to this combined dataframe, and used it as the source to impute the missing markers for panel 1. This process was repeated for all three panels. We tested this with SOM grid sizes of 4×4, 8×8, 12×12, and 16×16.
● **Evaluation:** Imputation accuracy was quantified using Spearman’s rank correlation between the imputed and original held-out marker values for each cell.

**Imputation on Synthetic Data:** To directly compare single-cell imputation accuracy, we used our synthetic dataset with known ground-truth values.

● **cyCombine Application:** We followed the same imputation strategy used for public datasets, with a 8×8 SOM grid (default).
● **UVAE Application:** We used the best-performing supervised model, and two unsupervised models (with or without MMD) from the synthetic benchmark.
● **Evaluation:** For both methods, we calculated the imputation MSE between the generated values and the known ground-truth values for the missing markers. We also computed EMD, and MAD scores, and re-computed iLISI and cLISI in the reconstructed data space (as opposed to latent space) to enable direct comparison with cyCombine.

#### COVID-19 data pre-processing and multi-stage integration strategy

The integration of the CIRCO COVID-19 clinical dataset involved a multi-stage pipeline, including initial data processing and cleaning, followed by a sequential integration strategy to harmonise the numerous distinct marker sets.

**Flow Cytometry Data Pre-processing:** Raw FCS files were processed using a custom R^30^ pipeline. Samples were first manually compensated for channel spillover and annotated in *FlowJo*^31^, and imported using the *CytoML*^32^ and *flowWorkspace*^33^ packages. A *Logicle* transform^34^ with default parameters was applied to marker values using *flowCore*^35^. We then used *openCyto*^36^ for semi-automated gating to identify and annotate doublets (*singletGate*) and debris (a two-stage *flowClust* gate on FSC/SSC and SSC/CD45). To harmonise feature names across different panel versions, we created a mapping of all observed marker names to a common, standardised set. For batch annotation, we used a concatenation of the sample acquisition date and processing date, ensuring that only samples acquired and processed on the same day were considered part of the same batch. From each processed file, a random subsample of 10,000 cells was taken for model training. To improve the training of classifiers on infrequent populations, an additional 1,000 cells from each manually defined cell type in the lineage panel were included during the training phase but were excluded from all downstream analyses. The final processed data was stored in an HDF5^37^ structure for use with the Python-based UVAE framework.

**Multi-Stage Integration Strategy:** Due to the large number of distinct marker sets (12 for lineage, 18 for chemokine), we adopted a multi-stage integration strategy to ensure stability and efficiency.

1. **Primary Integration:** We first identified the primary marker sets that covered the majority of the data (three sets for lineage, four for chemokine). A UVAE model was trained to integrate these primary sets in parallel. This model included a VAE for each primary set, with batch effect correction (conditioning and latent normalisation) applied within each. The latent spaces were merged using both a Subspace constraint on their shared markers and an MMD constraint applied between all pairs. Two shared classifiers were trained on the integrated latent space to predict cell types and doublets. Resampling for the lineage panel was guided by the cell-type classifier. For the chemokine panel, which is dominated by neutrophils, we used a hybrid resampling approach, guided by both the cell-type classifier and an unsupervised GMM clustering of each batch. Hyper-parameters for this primary model were selected via Bayesian optimisation.
2. **Sequential Integration:** After the primary integration model was trained and its weights frozen, we sequentially integrated the remaining, smaller marker sets. For each new set, an additional VAE was added to the model. Its latent space was aligned to the primary model by applying a Subspace constraint with the most similar primary marker set (based on feature overlap) and an MMD constraint with all primary sets. This new VAE was trained using the same optimal hyper-parameters identified during the primary integration phase, projecting the new data onto the stable, pre-existing latent space.
3. **Data Regeneration:** Once all marker sets were integrated, we generated a uniform, harmonised dataset. We defined a marker superset containing all features from the primary marker sets. For each cell, we used its specific encoder to obtain a latent representation and then used the decoders from the primary VAEs to reconstruct the full marker superset, correcting for batch effects by translating all cells to a single, arbitrarily chosen target batch.

#### Longitudinal regression models for predicting peak COVID severity

To assess the predictive value of the integrated COVID-19 data, we developed and evaluated several deep learning models for predicting a patient’s peak clinical severity from their longitudinal time-series of flow cytometry samples.

**Cross-Validation Strategy:** We implemented a 4-fold cross-validation scheme at the patient level. Patient IDs were randomly split into four folds. For each fold, which served as the held-out test set, a complete UVAE integration model was trained from scratch using only the data from the remaining three folds. This was done by first training a primary integration model in parallel on the largest marker sets from the training folds, then sequentially projecting the remaining training-fold marker sets onto it. For this process, we used a fixed set of hyper-parameters (*pull*=5, *frequency*=2 for MMD and Subspace constraints, *ease_epochs*=5, *lr_merge*=2) based on our experience with the main model. After training, the samples from the held-out test PIDs were treated as new, unseen panels and projected onto the trained model to generate harmonised data. This procedure prevents any information from the test set from influencing the batch correction or original latent space structure. This process also creates four different datasets (including both train and test folds). The regression models for each fold were trained on their corresponding training data and used to predict over the held-out data. The final test results were obtained by concatenating the predictions from the four held-out folds.

**Clustering for Proportion Models:** To avoid data leakage through clustering, neutrophil clusters for the proportion-based models were defined using only the training data for each fold. Neutrophils from the integrated training data were clustered using GMM (5 clusters) or Leiden (variable number of clusters). A K-Nearest Neighbours classifier (*k*=15) was then trained on these cluster assignments. This classifier was subsequently used to assign cluster labels to the neutrophils from the held-out test data.

**Regression Models and Inputs:** We encoded the peak severity categories (*Healthy*, *Mild*, *Moderate*, *Severe*) as ordinal regression targets (0.5, 2.0, 3.0, 4.5, respectively) and optimised models for MSE. For calculating classification scores, the predictions were rounded to the nearest integer, and compared to mapping between 1 and 4. The expansion of range for the lowest and highest category ensures that all four categories target approximately the same span of values (that is, *Healthy* samples can be predicted between 0.5 and 1.5, *Mild* between 1.5 and 2.5, etc.), which improves our accuracy for the *Healthy* and *Severe* categories. To enable comparison between models which use one or both panel inputs, we only used acute samples which contained both types of panel, resulting in 26 controls, and 41 mild, 47 moderate, 34 severe patients.

We tested two main classes of input data derived from the integrated whole blood (WB) and chemokine (Chemo) panels.

1. **Proportion-based Models:** These models used vectors of cell-type proportions (from the WB panel) and/or cluster proportions (for neutrophils in the Chemo panel) as input for each time-point. In linear models, the input was directly followed by a single fully-interconnected neuron with linear activation, applied independently to all non-zero inputs. The prediction was then max-pooled across all time-points to result in peak severity. In MLP models, a time-distributed dense layer with 50 neurons and *ReLU* activation was independently applied to each time-point. Consecutively, the max-pooling models applied a single linear neuron, followed by global maximum. The RNN pooling models applied an RNN layer with 50 units and *tanh* activation, followed by a single linear neuron.
2. **Set-based Models:** These models operated directly on the raw marker values. For each time-point, a random set of 20 or 100 cells was sampled for each included panel type. This set was processed by a permutation-invariant network arm consisting of a 1D convolution (50 filters of size 1, with *ReLU* activation), followed by parallel global max– and average-pooling, outputs of which were concatenated. The output of this network arm is invariant to the permutation of cells given as input. A separate arm was trained for WB and chemokine inputs, and if both were used, their outputs were concatenated for each time-point. For a max-pooling model, an additional time-distributed dense layer with 50 neurons and *ReLU* activation was applied before a single linear neuron and global max-pooling. The RNN model did not use the extra dense layer but instead applied an RNN layer (with 50 units and *tanh* activation), followed by a linear neuron.

**Model Training and Evaluation:** For the cross-validation, regression models were trained on two of the three non-test folds, with the third used as a validation set for early stopping. After three models were obtained, they were used in an ensemble to predict on the held-out fold, with their outputs averaged. All models were trained for a maximum of 2000 epochs with early stopping. To handle the class imbalance in severity groups, samples were up-sampled with replacement during training. For set-based models, a new random set of cells was sampled for each training sample per epoch. To predict a validation loss, this sampling was repeated 10 times, with the losses averaged. For final prediction, the sampling was repeated 100 times for each sample, with the predictions averaged. Performance was evaluated using test MSE and F1 score.

For the trajectory visualizations (Figure 7) and gradient attribution plots (Figure S10), a single UVAE model trained on the globally integrated dataset (all data included) was used to generate the data for set model training. Max-pooling model with 20-cells input from both WB and Chemokine panels was used for trajectory predictions, and a similar RNN-pooling model with only Chemokine input was used for gradient computation.

### QUANTIFICATION AND STATISTICAL ANALYSIS

All statistical analyses were performed using Python (v3.10.4) with the scipy.stats^26^ and statsmodels packages^27^ or R (v4.2.0). Number of patients and patient-timepoint samples are given in Table 6, S3 and S4. Randomization for cross-validation was performed at the patient ID level.

For comparisons of cell-type proportions and marker levels between patient groups and healthy controls, statistical significance was assessed using Welch’s t-test, which does not assume equal variance. The samples used for these tests represent individual patient time-points (*n* is the number of patient time-point samples in each group). For cell-type analysis, the input value for each sample was the proportion of a given cell type relative to all live, single cells. For marker analysis, the input value was the mean expression level of that marker across all cells of a given type (e.g., neutrophils) within the sample.

To correct for multiple hypothesis testing, the raw *p*-values for each family of tests (e.g., all cell-type proportion comparisons in acute samples) were adjusted using the Benjamini-Hochberg procedure to control the false discovery rate. A family of tests consisted of comparisons between healthy controls and each of the three severity subgroups (Mild, Moderate, Severe), as well as a comparison between controls and all patient severities combined. An adjusted *p*-value (*q*-value) < 0.05 was considered statistically significant.

Performance metrics for data integration benchmarks included Earth Mover’s Distance (EMD), Mean Absolute Deviation (MAD), Mean Squared Error (MSE), Spearman’s rank correlation, and Local Inverse Simpson’s Index (LISI). LISI scores on the synthetic data benchmark were calculated with a perplexity of 100 using the implementation from the Harmony package^24^.

Performance of the longitudinal regression models was evaluated using test set MSE and macro-averaged F1 scores.

In all boxplots, the centre line indicates the median, the whiskers begin at the first and third quartiles and extend to the furthest data point within 1.5 times the interquartile range (IQR) from their start. Data points beyond the whiskers are considered outliers.

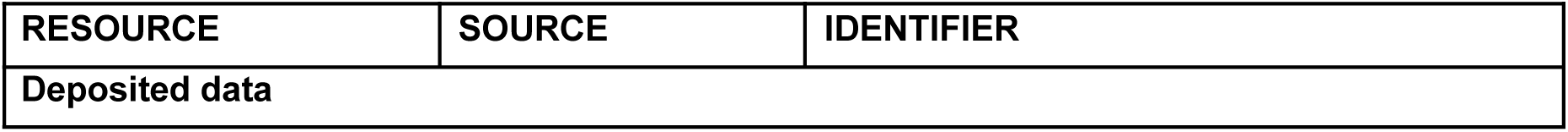
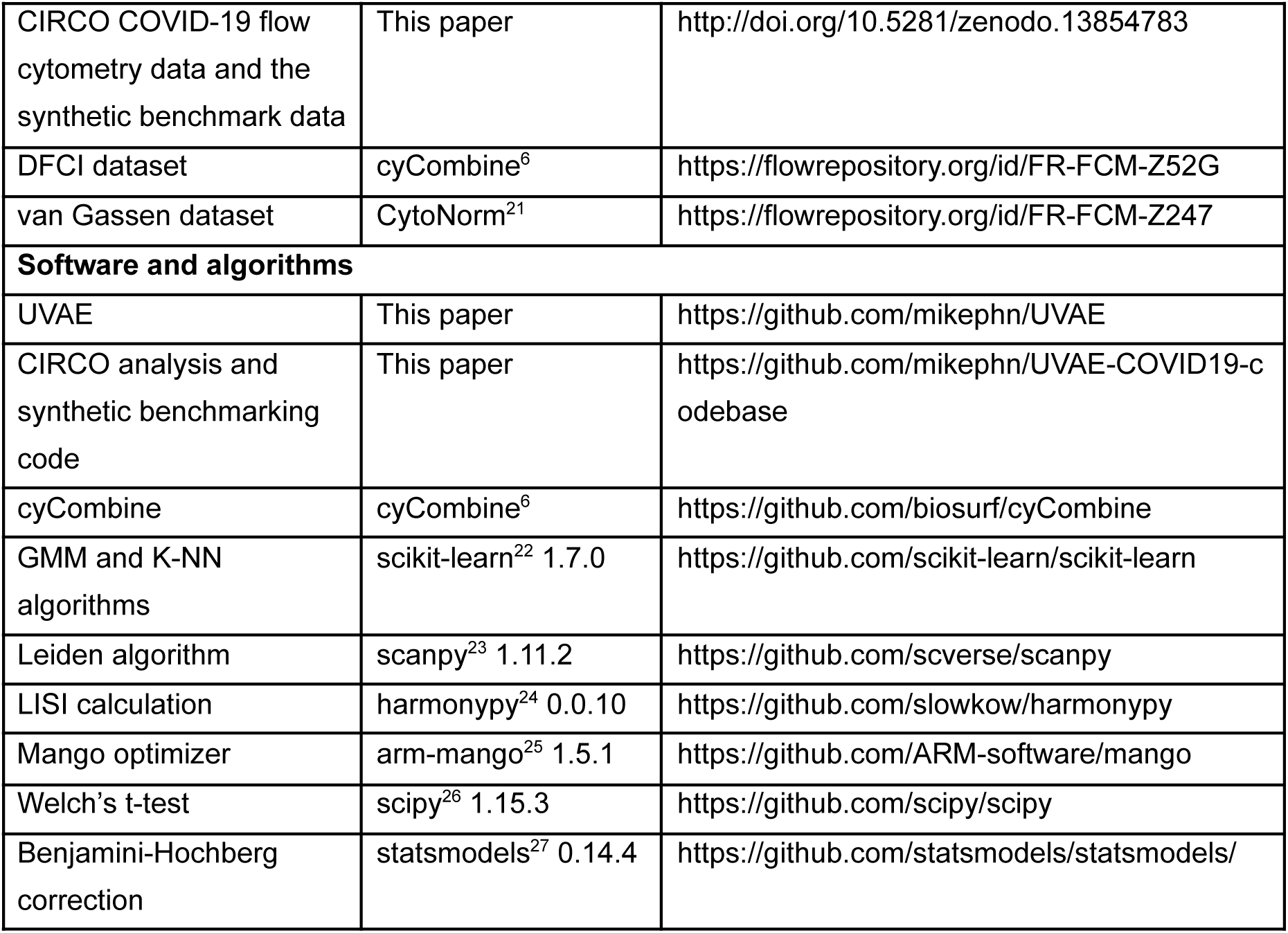
KEY RESOURCES TABLE.

## Notes

### Competing Interest Statement

The authors have declared no competing interest.

### Summary of Updates

Additional benchmarks were added. Manuscript was re-written for better readability.

