## Supplementary for "Integration of Unpaired and Heterogeneous Clinical Flow Cytometry Data"

**Algorithm S1** UVAE training with multiple constraints and batch correction.
Related to STAR Methods: *UVAE Framework: General Overview*.

**Require:** Data $D$, labels $Y$, conditions $B$, learning rate $\eta$

Initialize constraints and their hyperparameters; index their corresponding data:
  Autoencoders ($A_p$), Classifier ($C$), Subspace ($S, pull_S$), MMD ($M, pull_M$), Normalization ($N$)
Define resampling links: $C \Rightarrow N$ and $C \Rightarrow M$
**for** epoch $ep = 1, \ldots, E$ **do**
  $O \Leftarrow$ ShuffledTrainingOrder($\mathcal{C}_{\text{trainable}}$) ▷ Create shuffled order of constraints to train
  **for** $const$ in $O$ **do**
    $m \Leftarrow$ SampleMinibatch($const$, batch_size)
    Let $(x_m, b_m)$ be the data and conditions for indexes $m$
    $z, z_{\text{mean}}, z_{\text{log\_var}} \Leftarrow enc_{p \ni m}(x_m, b_m)$ ▷ Encode minibatch into latent space
    **if** $const \in \{A_p\}$ **then** ▷ Case 1: Autoencoder reconstruction
      $L_{AE} \Leftarrow \text{MSE}(dec_p(z, b_m), x_m) + \beta \cdot \text{KL}(z_{\text{mean}}, z_{\text{log\_var}})$
      $\theta_{A_p} \Leftarrow \theta_{A_p} - \eta \nabla_{\theta_{A_p}} L_{AE}$
    **else** ▷ Case 2: Supervised or alignment constraint
      $z_n \Leftarrow z - \text{offsets}(enc_{p \ni m}, b_m)$ ▷ Apply latent normalization
      $L_{const}, L_{merge} \Leftarrow const(m, z_n)$ ▷ Calculate primary and merging losses
      **if** $L_{const} \neq 0$ **then**
        $\theta_{const} \Leftarrow \theta_{const} - \eta \nabla_{\theta_{const}} L_{const}$ ▷ Update constraint parameters (e.g., Classifier)
      **end if**
      **if** $L_{merge} \neq 0$ **then**
        $\theta_{enc} \Leftarrow \theta_{enc} - \eta \nabla_{\theta_{enc}} (pull_{const} \cdot L_{merge})$ ▷ Update encoders to align latent space
      **end if**
    **end if**
  **end for**
  $prop_{ease} \Leftarrow \min(1, ep/\text{ease\_epochs})$
  Update resampling indexes for Normalization ($N$) and MMD ($M$) based on Classifier ($C$) predictions, using $prop_{ease}$.
  Update normalization offsets in all encoders using Normalization constraint ($N$) and the newly resampled indexes, with $prop_{ease}$.
**end for**

**Algorithm S2** Batch of classifier training.
Related to STAR Methods: *UVAE Framework: General Overview*.

**Require:** Minibatch indexes $m$, normalized input embedding $z_n$, learning rate $\eta$
**Require:** Initialized classifier function $func_C(z)$ with parameters $\theta_C$
**Require:** One-hot targets $y$ for minibatch $m$
$y' \Leftarrow func_C(z_n)$ ▷ Predict targets for the minibatch
$L_C \Leftarrow$ CategoricalCrossentropy$(y, y')$ ▷ Calculate classification error
$\theta_C \Leftarrow \theta_C - \eta\nabla_{\theta_C} L_C$ ▷ Update classifier parameters to minimize error
$L_{merge} \Leftarrow L_C$ ▷ Assign classification loss as the merging loss for encoders

**Algorithm S3** Calculating latent normalization offsets.
Related to STAR Methods: *Batch Effect Correction Mechanisms*.

**Require:** Normalization constraint $N$ with batch assignments $B$, an encoder $enc$
**Require:** Balanced/resampled data indexes $idx_{res}$
**Require:** Easing-in proportion $prop$
$Z \Leftarrow enc(D(idx_{res}), B(idx_{res}))$ ▷ Embed the resampled data into latent space
$\mu \Leftarrow \varnothing$ ▷ Initialize map to store mean embedding for each batch
**for** $b$ in unique$(B(idx_{res}))$ **do**
    $Z_b \Leftarrow Z[B(idx_{res}) == b]$ ▷ Get latent vectors corresponding to batch $b$
    **if** len$(Z_b) > 0$ **then**
        $\mu_b \Leftarrow$ mean$(Z_b)$ ▷ Calculate and store the mean embedding
    **end if**
**end for**
**if** a target batch $b_t$ is defined in $N$ **then**
    $\mu_{target} \Leftarrow \mu_{b_t}$ ▷ Set target to a specific batch's mean
**else**
    $\mu_{target} \Leftarrow$ mean(values$(\mu)$) ▷ Set target to the global mean of all batch means
**end if**
**for** $b$ in keys$(\mu)$ **do**
    $\delta_b \Leftarrow \mu_b - \mu_{target}$ ▷ Calculate the offset vector from the target
    $offsets_{enc,b} \Leftarrow \delta_b \cdot prop$ ▷ Update the stored offset for this encoder-batch pair
**end for**

**Algorithm S4** Batch of panel merging with a shared channel subspace.
Related to STAR Methods: *Integration of Disparate Panels*.

**Require:** Minibatch indexes $m$, normalized panel embedding $z_n$, learning rate $\eta$
**Require:** Initialized Subspace autoencoder $S$ with encoder $enc_S$, decoder $dec_S$, and parameters $\theta_S$
**Require:** Subspace pull strength $pull_S$
$(x_m, b_m) \Leftarrow$ get_data_and_conditions$(m)$
$x_{sh} \Leftarrow x_m[:,$ shared_channels$]$ ▷ Subset input to only shared channels
$z_S \Leftarrow enc_S(x_{sh}, b_m)$ ▷ Embed shared channels with the Subspace encoder
$x'_{sh} \Leftarrow dec_S(z_S, b_m)$ ▷ Reconstruct shared channels
$L_{S\_rec} \Leftarrow$ MSE$(x'_{sh}, x_{sh})$ ▷ Subspace's internal reconstruction loss
$z_{Sn} \Leftarrow z_S -$ offsets$(enc_S, b_m)$ ▷ Apply normalization to the Subspace embedding
$L_{merge} \Leftarrow$ MSE$(z_{Sn}, z_n)$ ▷ Calculate merging loss between Subspace and panel embeddings
$\theta_S \Leftarrow \theta_S - \eta\nabla_{\theta_S}(L_{S\_rec} + pull_S \cdot L_{merge})$ ▷ Update Subspace AE on combined loss

**Algorithm S5** Batch of panel merging with MMD.
Related to STAR Methods: *Integration of Disparate Panels*.

**Require:** Minibatch indexes $m$ and their normalized embedding $z_n$
**Require:** Condition assignments $B$ for indexes $m$, sampled to contain an equal number of samples, $n$, from exactly two distinct conditions, $b_1$ and $b_2$.

$z_1 \Leftarrow z_n[B == b_1]$ ▷ Get embeddings for the first condition
$z_2 \Leftarrow z_n[B == b_2]$ ▷ Get embeddings for the second condition
$n \Leftarrow \text{len}(z_1)$
Let $\Gamma$ be a predefined set of scale parameters for the RBF kernel.
Define multiscale RBF kernel $k(z, z') \Leftarrow \sum_{\gamma \in \Gamma} \exp(-\gamma \|z - z'\|^2)$.
$L_{merge} \Leftarrow \frac{1}{n^2} \sum_{i,j=1}^{n} (k(z_{1i}, z_{1j}) + k(z_{2i}, z_{2j}) - 2k(z_{1i}, z_{2j}))$ ▷ Empirical MMD loss

**Algorithm S6** Supervised resampling of a constraint.
Related to STAR Methods: *Class-Balanced Resampling*.

**Require:** Classifier $C$ with function $func_C$, target constraint $T$ to be resampled
**Require:** All data indexes $idx_{all}$ for constraint $T$, with batch assignments $B$
**Require:** Resampling proportion $prop$, boolean $dropMissing$

$idx_{res} \Leftarrow \varnothing$ ▷ Initialize empty set for resampled indexes
$Z \Leftarrow enc(D(idx_{all}), B)$ ▷ Embed all data covered by the target constraint
$Z_n \Leftarrow Z - \text{offsets}(enc, B)$ ▷ Apply latent normalization
$Y_{pred} \Leftarrow func_C(Z_n)$ ▷ Predict classes for all samples
**for** $b$ in unique$(B)$ **do** ▷ First, count class instances in each batch
  **for** $c$ in unique$(Y_{pred})$ **do**
    $N_{bc} \Leftarrow \text{count}((B == b) \wedge (Y_{pred} == c))$
  **end for**
**end for**
**if** $dropMissing$ **then** ▷ Optionally ignore classes not present in all batches
  **for** $c$ in unique$(Y_{pred})$ **do**
    **if** $\exists b : N_{bc} == 0$ **then**
      $N_{bc} \Leftarrow 0$ for all $b$ ▷ Invalidate class c if missing anywhere
    **end if**
  **end for**
**end if**
$N_{total} \Leftarrow \sum_{b,c} N_{bc}$
**for** $c$ in unique$(Y_{pred})$ **do** ▷ Calculate average proportion of each class
  $P_c \Leftarrow (\sum_b N_{bc})/N_{total}$
**end for**
**for** $b$ in unique$(B)$ **do** ▷ Resample each batch to match average proportions
  $N_b \Leftarrow \text{count}(B == b)$
  **for** $c$ in unique$(Y_{pred})$ **do**
    $idx_{bc} \Leftarrow \text{get\_indexes}((B == b) \wedge (Y_{pred} == c))$
    $N_{to\_sample} \Leftarrow \text{round}(prop \cdot P_c \cdot N_b)$
    $idx_{sampled} \Leftarrow \text{RandomSample}(idx_{bc}, N_{to\_sample})$ ▷ Sample with replacement
    $idx_{res} \Leftarrow idx_{res} \cup idx_{sampled}$
  **end for**
**end for**
$idx_{unbalanced} \Leftarrow \text{RandomSample}(idx_{all}, \text{round}((1 - prop) \cdot \text{len}(idx_{all})))$
$idx_{res} \Leftarrow idx_{res} \cup idx_{unbalanced}$ ▷ Fill remainder with random unbalanced samples

**Algorithm S7** Unsupervised resampling with external batch-level clustering.
Related to STAR Methods: *Class-Balanced Resampling*.

**Require:** Target constraint $T$ to be resampled (e.g., Normalization)
**Require:** All data indexes $idx_{all}$ for constraint $T$
**Require:** Batch assignments $B$ for indexes $idx_{all}$
**Require:** External cluster assignments $C_{ext}$ for indexes $idx_{all}$
**Require:** Resampling proportion $prop$

$idx_{res} \Leftarrow \varnothing$ ▷ Initialize empty set for balanced indexes
**for** $b$ in unique($B$) **do** ▷ Iterate through each batch independently
  $idx_b \Leftarrow$ get_indexes($B == b$) ▷ Get all indexes for the current batch
  Let $C_{ext\_b}$ be the cluster assignments for indexes $idx_b$
  $N_b \Leftarrow$ len($idx_b$) ▷ Total number of samples in this batch
  $K_b \Leftarrow$ len(unique($C_{ext\_b}$)) ▷ Number of clusters in this batch
  **if** $K_b > 0$ **then**
    $N_{per\_cluster} \Leftarrow$ round($(prop \cdot N_b)/K_b$) ▷ Calculate sample count per cluster
    **for** $k$ in unique($C_{ext\_b}$) **do** ▷ For each cluster within the batch
      $idx_{bk} \Leftarrow$ get_indexes($(B == b) \wedge (C_{ext} == k)$)
      $idx_{sampled} \Leftarrow$ RandomSample($idx_{bk}, N_{per\_cluster}$) ▷ Sample with replacement
      $idx_{res} \Leftarrow idx_{res} \cup idx_{sampled}$ ▷ Add to the balanced index set
    **end for**
  **end if**
**end for**
$idx_{unbalanced} \Leftarrow$ RandomSample($idx_{all}$, round($(1 - prop) \cdot$ len($idx_{all}$)))
$idx_{res} \Leftarrow idx_{res} \cup idx_{unbalanced}$ ▷ Fill remainder with random unbalanced samples

**Algorithm S8** Imputation of a missing channel.
Related to STAR Methods: *Generation of Homogenized Data*.

**Require:** Sample indexes $m$ from a source panel $s$
**Require:** The name of the missing channel, $ch_{miss}$
**Require:** The full set of autoencoders $\{A_p\}$, encoders $\{enc_p\}$, and decoders $\{dec_p\}$
**Require:** All batch assignments $B$ and the learned normalization offsets
**Require:** Optional user-provided target conditions $B_{target}$

$(x_s, b_s) \Leftarrow$ get_data_and_conditions($m$, from source panel $s$)
$z_s \Leftarrow enc_s(x_s, b_s)$ ▷ Embed source data into latent space
$z_n \Leftarrow z_s -$ offsets($enc_s, b_s$) ▷ Apply source normalization to get the unified embedding
$X'_{miss} \Leftarrow$ empty_array ▷ Initialize accumulator for imputed channel values
**for** each panel $p$ in the model **do**
  **if** $ch_{miss}$ exists in the channels of panel $p$ **then**
    **if** a valid target condition $b_t$ for panel $p$ is provided in $B_{target}$ **then**
      $b_{target} \Leftarrow b_t$ ▷ Use user-specified target condition
    **else**
      $b_{target} \Leftarrow$ get_first_valid_condition($p$) ▷ Default to first valid condition for decoder $p$
    **end if**
    $z_{rebiased} \Leftarrow z_n +$ offsets($enc_p, b_{target}$) ▷ Re-apply the target condition's offset
    $x'_p \Leftarrow dec_p(z_{rebiased}, b_{target})$ ▷ Decode using the conditional decoder and re-biased latent
    $x'_{pch} \Leftarrow x'_p[:, ch_{miss}]$ ▷ Extract the values for the imputed channel
    $X'_{miss} \Leftarrow$ append_column($X'_{miss}, x'_{pch}$) ▷ Collect predictions from this decoder
  **end if**
**end for**
$x'_{avg} \Leftarrow$ mean($X'_{miss}$, axis=1) ▷ Average reconstructions from all relevant decoders

Table S1: Best performing combinations of batch correction and merging constraints, with or without resampling on Whole Blood synthetic dataset, related to Table 2.

| Configuration[a] | | | | | Metrics | | | | | | |
|---|---|---|---|---|---|---|---|---|---|---|---|
| | | | | | Overall | MSE imputed | | iLISI | | cLISI | |
| Conditioning[b] | Norm | MMD[c] | Subspace | Resampling[d] | rank | median | rank | median | rank | median | rank |
| ✓ | ✓ | ✓ | ✓ | ✓ | **1** | .31 | **4** | 6.48 | **2** | 1.27 | **2** |
| | ✓ | ✓ | ✓ | ✓ | **2** | .31 | **5** | 6.61 | **1** | 1.28 | **4** |
| | ✓ | | ✓ | ✓ | **3** | .29 | **1** | 5.65 | **5** | 1.29 | 8 |
| | | ✓ | ✓ | ✓ | **4** | .33 | 9 | 5.96 | **3** | 1.28 | 6 |
| ✓ | | ✓ | ✓ | ✓ | **5** | .32 | 6 | 5.69 | **4** | 1.3 | 12 |
| ✓ | ✓ | | ✓ | ✓ | 6 | .29 | **2** | 4.59 | 16 | 1.28 | **5** |
| | ✓ | ✓ | ✓ | | 7 | .32 | 7 | 5.5 | 8 | 1.3 | 11 |
| | ✓ | ✓ | | ✓ | 8 | .36 | 16 | 5.28 | 11 | 1.26 | **1** |
| ✓ | | ✓ | ✓ | | 9 | .34 | 13 | 5.32 | 9 | 1.29 | 7 |
| ✓ | | ✓ | | ✓ | 10 | .35 | 15 | 5.55 | 6 | 1.29 | 9 |
| ✓ | ✓ | ✓ | ✓ | | 11 | .33 | 12 | 4.81 | 15 | 1.27 | **3** |
| ✓ | | | ✓ | | 12 | .29 | **3** | 3.93 | 21 | 1.3 | 13 |
| | ✓ | | ✓ | | 13 | .33 | 10 | 5.17 | 14 | 1.3 | 14 |
| | | ✓ | | ✓ | 14 | .38 | 17 | 5.26 | 12 | 1.29 | 10 |
| | | ✓ | ✓ | | 15 | .33 | 11 | 5.2 | 13 | 1.32 | 16 |
| | | | ✓ | | 16 | .35 | 14 | 5.31 | 10 | 1.32 | 17 |
| ✓ | ✓ | ✓ | | ✓ | 17 | .4 | 18 | 5.54 | 7 | 1.4 | 21 |
| ✓ | ✓ | | ✓ | | 18 | .32 | 8 | 3.38 | 22 | 1.33 | 19 |
| | ✓ | ✓ | | | 19 | .42 | 19 | 4.15 | 20 | 1.31 | 15 |
| ✓ | ✓ | ✓ | | | 20 | .42 | 20 | 4.35 | 18 | 1.33 | 18 |
| | | ✓ | | | 21 | .42 | 21 | 4.44 | 17 | 1.33 | 20 |
| ✓ | | ✓ | | | 22 | .43 | 22 | 4.27 | 19 | 1.43 | 22 |

[a]Shared settings: VAE (50 dim, 2 hidden layers, 256 neurons each), unsupervised LISI target, classifier fitted.
[b]Conditioning both encoder and decoder.
[c]MMD between panels only.
[d]Resampling normalisation/MMD using manual cell-type annotation.

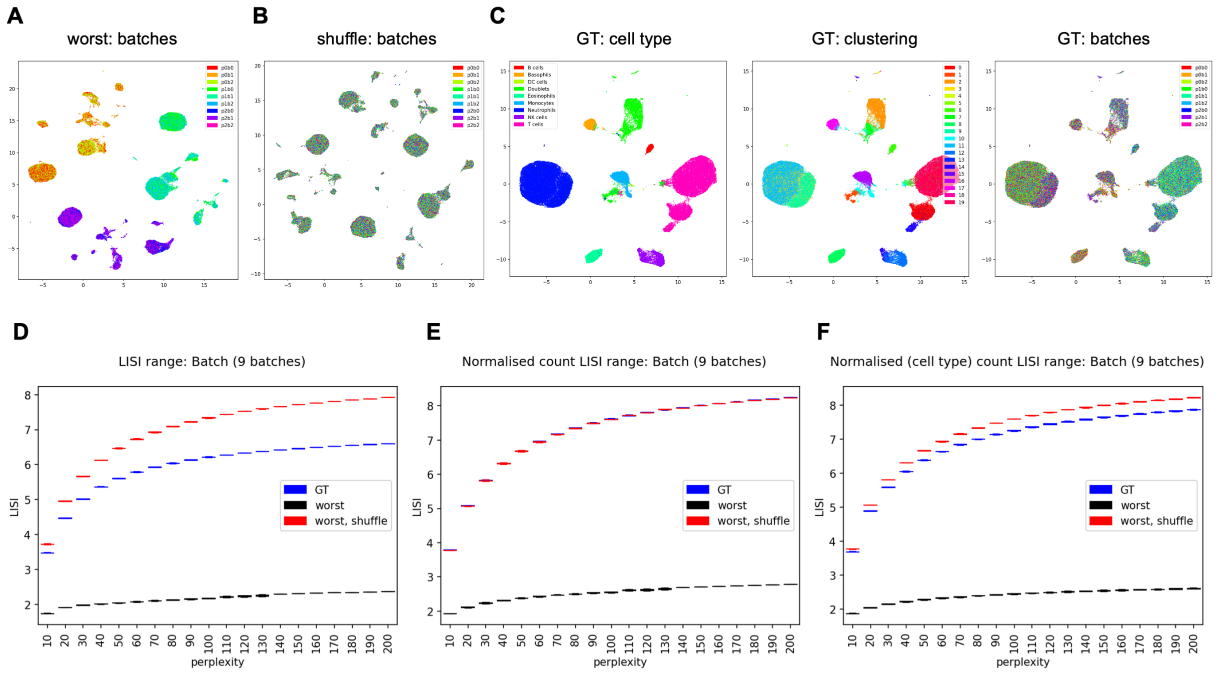


Figure S1: **Normalised LISI Scoring**. Related to STAR Methods: *Model Selection and Hyperparameter Optimisation*. (A) UMAP of VAE embedding of the synthetic dataset without batch correction or merging applied, showcasing worst-case model with maximum expected batch separation. (B) Expected worst case of uninformative over-alignment, created by randomly shuffling batch labels. (D) UMAP of VAE embedding of the ground-truth data without panel splitting or batch effects, showing correct cell-type, clustering, and batch alignment. Note that optimal batch alignment is not perfectly uniform. (D)-(F) LISI scores calculated for different perplexities using ground truth model (GT), worst case model of maximum separation (worst), and worst case model of maximum over-mixing (worst, shuffle). Medians across all cells were calculated over 10 repeated model trainings for each configuration. (D) Unnormalised LISI scores calculated without accounting for class imbalance between batches. Shuffled model scores higher than GT. (E) Normalised LISI scores calculated by down-sampling each ground truth (GT) cluster in each batch to the lowest count across all batches. GT model scores the same as shuffled, but GT class assignment would typically be unknown. (F) Normalised LISI scores calculated by down-sampling each manually annotated cell-type in each batch to the lowest count across all batches. Shuffled model has smaller advantage over GT than without this normalization.

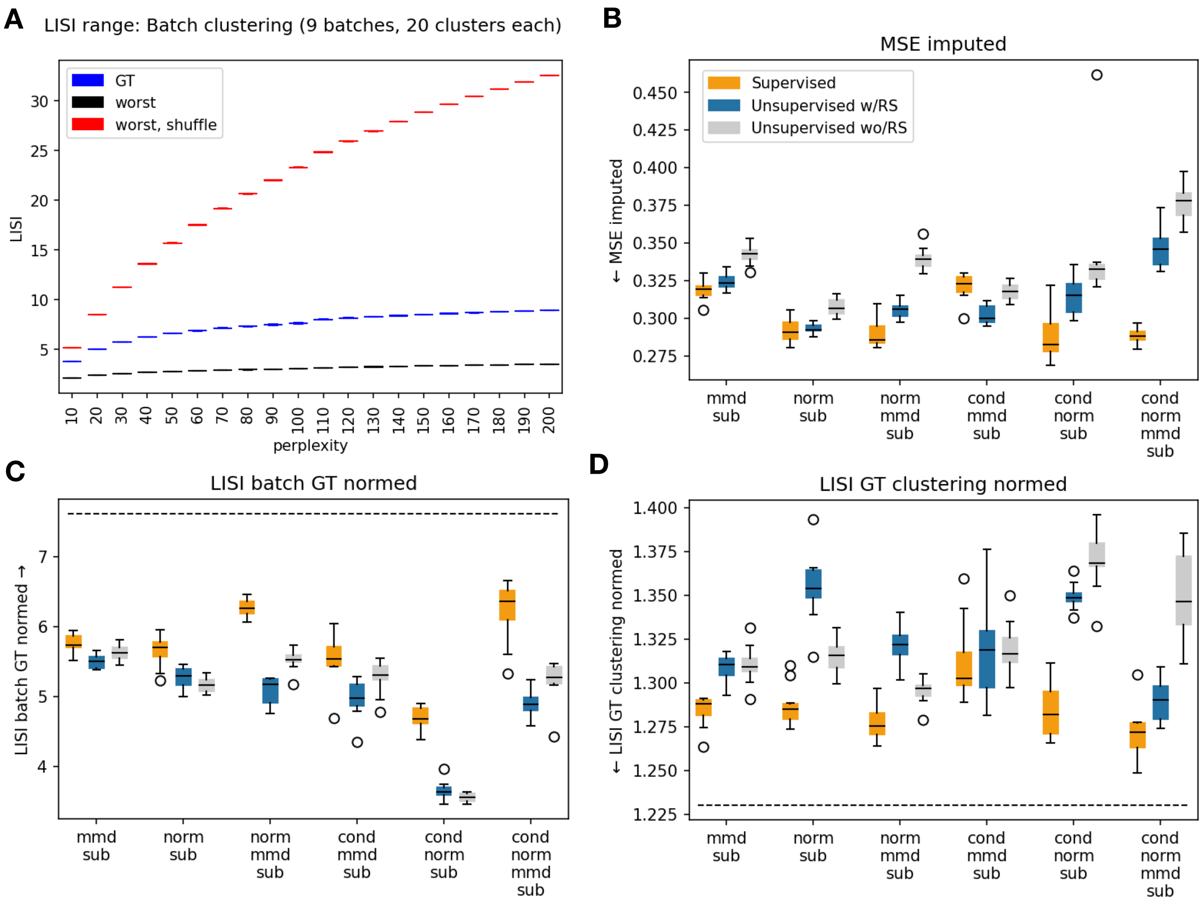


Figure S2: **Unsupervised LISI Scoring and Resampling**. Related to STAR Methods: *Model Selection and Hyperparameter Optimisation*. (A) LISI scores on Whole Blood synthetic dataset calculated using independent clustering of 9 overlapping batches to 20 clusters each. Scores were calculated for different perplexities using ground truth model (GT), worst case model of maximum separation (worst), and worst case model of maximum over-mixing (worst, shuffle). Medians across all cells were calculated over 10 repeated model trainings. (B)-(D) Imputation and alignment comparison for supervised models (using predicted cell-type LISI optimisation target and cell-type resampling), or unsupervised models using independent clustering of each batch. Unsupervised w/RS indicates the batch clustering was used to balance latent normalisation (norm) or between-panel MMD. Unsupervised wo/RS indicates no resampling was performed. All models shared settings: VAE with 50 dim, 2 hidden layers, 256 neurons each. 'cond': encoder and decoder conditioning of batch. 'norm': latent space normalisation. 'sub': subspace merging. Dotted lines indicate scores obtained by ground-truth model trained on unaltered data. (B) Imputation mean-squared errors. (C) LISI scores of batch alignment normalised with ground-truth cluster split (higher is better). (D) Normalised LISI scores of ground-truth cluster separation (lower is better).

Table S2: Configurations with supervised (cell-type) or unsupervised (batch clustering) LISI optimisation target on Whole Blood synthetic dataset. Related to STAR Methods: *Model Selection and Hyperparameter Optimisation*

| Configuration[a] | | | | | | Metrics | | | | | |
|---|---|---|---|---|---|---|---|---|---|---|---|
| Conditioning[b] | Norm | MMD[c] | Subspace | LISI supervised[d] | Overall rank | MSE imputed median | MSE imputed rank | iLISI median | iLISI rank | cLISI median | cLISI rank |
| | ✓ | ✓ | ✓ | | **1** | .32 | **4** | 5.5 | **1** | 1.3 | **4** |
| | ✓ | ✓ | ✓ | ✓ | **2** | .33 | 7 | 5.46 | **3** | 1.3 | **5** |
| ✓ | | ✓ | ✓ | | **3** | .34 | 12 | 5.32 | **4** | 1.29 | **2** |
| | | | ✓ | ✓ | **4** | .32 | **3** | 5.26 | 7 | 1.31 | 10 |
| | ✓ | | ✓ | ✓ | **5** | .33 | 6 | 5.3 | 6 | 1.31 | 11 |
| ✓ | ✓ | ✓ | ✓ | | 6 | .33 | 10 | 4.81 | 13 | 1.27 | **1** |
| ✓ | | ✓ | ✓ | ✓ | 7 | .33 | 11 | 5.19 | 9 | 1.3 | 6 |
| | ✓ | | ✓ | | 8 | .33 | 8 | 5.17 | 10 | 1.3 | 8 |
| ✓ | ✓ | ✓ | ✓ | ✓ | 9 | .35 | 16 | 4.88 | 11 | 1.3 | **3** |
| ✓ | | | ✓ | | 10 | .29 | **1** | 3.93 | 22 | 1.3 | 7 |
| | | ✓ | ✓ | | 11 | .33 | 9 | 5.2 | 8 | 1.32 | 16 |
| | | ✓ | ✓ | ✓ | 12 | .35 | 14 | 5.46 | **2** | 1.32 | 17 |
| | | | ✓ | | 13 | .35 | 13 | 5.31 | **5** | 1.32 | 18 |
| ✓ | | | ✓ | ✓ | 14 | .3 | **2** | 3.95 | 21 | 1.31 | 13 |
| | ✓ | ✓ | | ✓ | 15 | .39 | 17 | 4.36 | 16 | 1.3 | 9 |

[a] Top 15 highest-scoring configurations shown. Shared settings: VAE (50 dim, 2 hidden layers, 256 neurons each), classifier fitted, no resampling.

[b] Conditioning both encoder and decoder.

[c] MMD between panels only.

[d] Supervised indicates cell-type annotation. Lack of tick mark indicates unsupervised batch clustering.

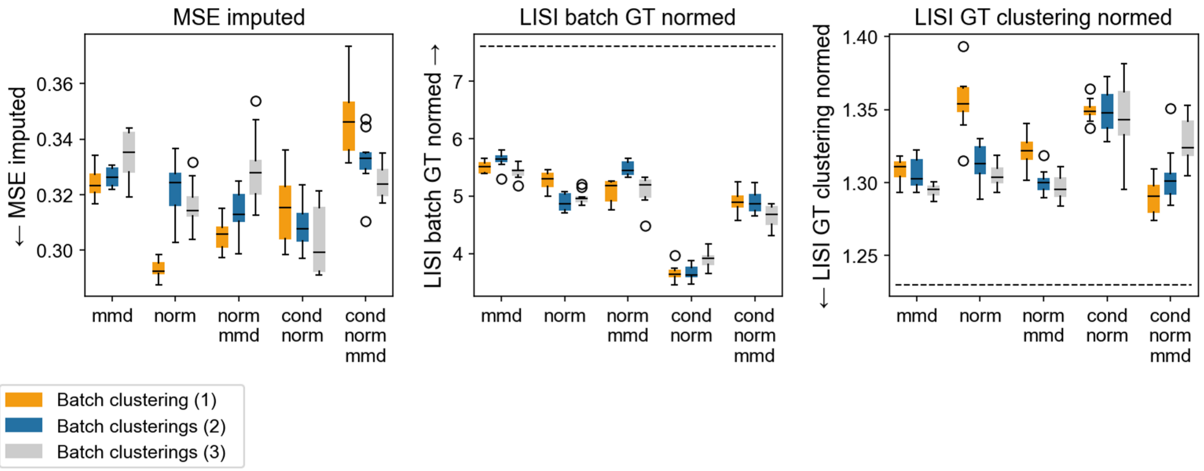


Figure S3: **Unsupervised Resampling with Multiple Cluster Assignments**. Related to STAR Methods: *Class-Balanced Resampling*. Imputation and alignment scores for unsupervised model configurations, where resampling was performed with increasing number of clusterings. One clustering used 20 clusters per batch, two used 20 and 15 clusters per batch, three used 20, 15 and 10 clusters per batch. Each clustering was used to balance an equal part of the data. All models shared settings: VAE with 50 dim, 2 hidden layers, 256 neurons each, subspace merging. 'cond': encoder and decoder conditioning of batch. 'norm': latent space normalisation. 'mmd': applied between panels. Dotted lines indicate scores obtained by ground-truth model trained on unaltered data.

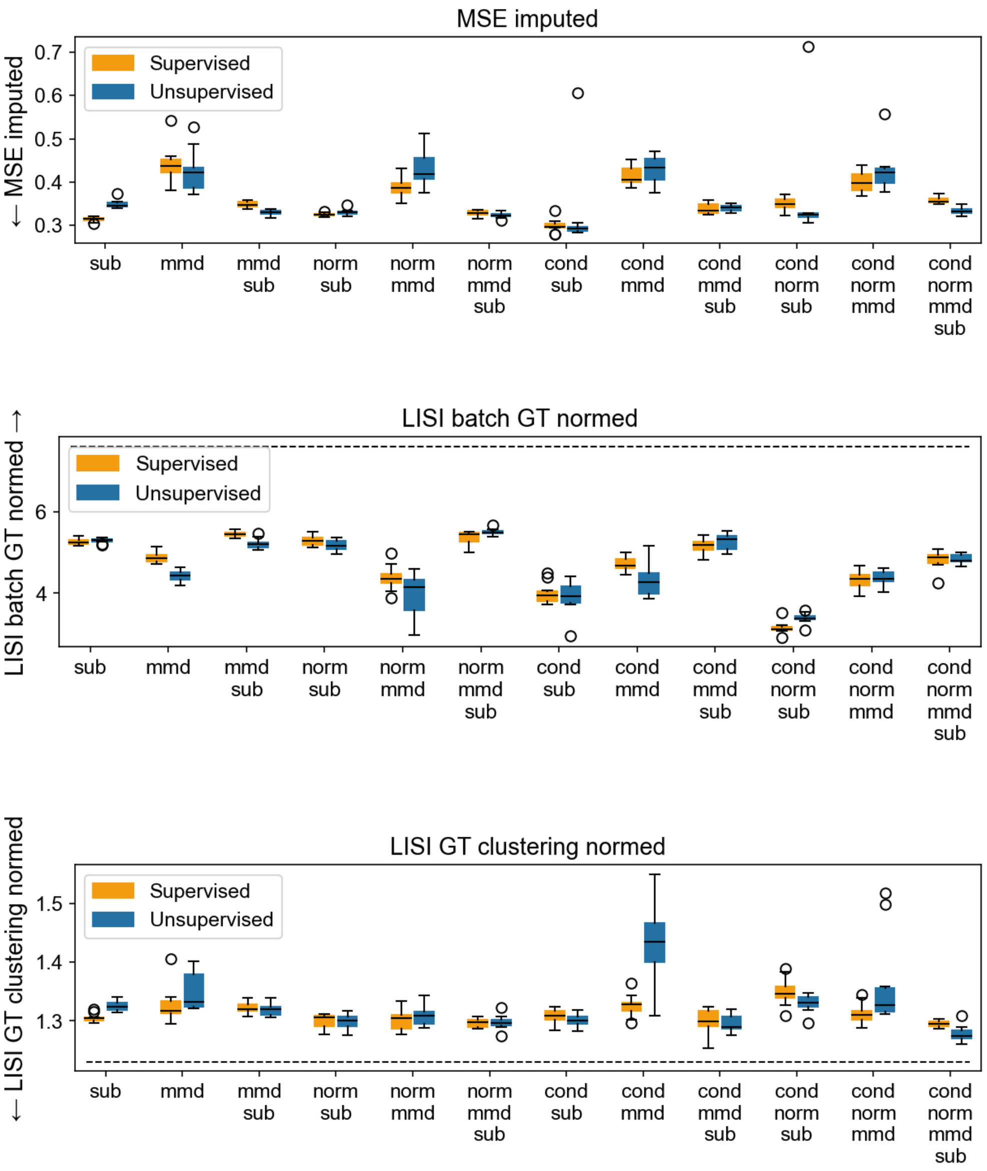


Figure S4: **Supervised vs. Unsupervised LISI Scoring without Resampling**. Related to STAR Methods: *Model Selection and Hyperparameter Optimisation*. Imputation and alignment scores for model configurations optimised using either supervised (cell-type prediction) or unsupervised (independent batch clustering) LISI metric. Models were otherwise kept identical and in both cases did include classifier training (necessary for supervised LISI) but did not use it for resampling during training. All models shared settings: VAE with 50 dim, 2 hidden layers, 256 neurons each. 'cond': encoder and decoder conditioning of batch. 'norm': latent space normalisation. 'mmd': applied between panels. 'sub': subspace merging. Dotted lines indicate scores obtained by ground-truth model trained on unaltered data.

Table S3: Significance of differences in cell-type proportions (Welch's t-test, Benjamini-Hochberg correction) between acute samples and controls. Cell-types predicted by the model are compared to manually labelled. Non-zero values refer to number of samples where at least one cell of given cell-type was labelled or predicted. P-values for rows labelled as Acute are calculated between controls and all acute samples pooled; for other rows they are between controls and a given severity only. Related to Figure 5.

| | **P-value** | | **Q-value** | | **Non-zero controls** | | **Non-zero acute** | |
|---|---|---|---|---|---|---|---|---|
| | Predicted | Labelled | Predicted | Labelled | Predicted | Labelled | Predicted | Labelled |
| **B cells** | | | | | | | | |
| Acute | 1.45E-01 | 2.54E-01 | 1.78E-01 | 3.02E-01 | 39 | 35 | 248 | 211 |
| Mild | 2.03E-01 | 7.90E-01 | 2.40E-01 | 8.15E-01 | 39 | 35 | 77 | 63 |
| Moderate | 6.70E-02 | 9.03E-02 | 8.57E-02 | 1.58E-01 | 39 | 35 | 90 | 75 |
| Severe | 4.85E-01 | 1.97E-01 | 5.54E-01 | 2.42E-01 | 39 | 35 | 81 | 73 |
| **Basophils** | | | | | | | | |
| Acute | 1.55E-10 | 9.35E-02 | 1.24E-09 | 1.58E-01 | 40 | 24 | 201 | 110 |
| Mild | 3.82E-09 | 1.52E-01 | 1.75E-08 | 2.22E-01 | 40 | 24 | 62 | 37 |
| Moderate | 7.38E-11 | 1.91E-01 | 7.87E-10 | 2.42E-01 | 40 | 24 | 79 | 38 |
| Severe | 2.02E-08 | 3.66E-02 | 6.45E-08 | 9.02E-02 | 40 | 24 | 60 | 35 |
| **DC cells** | | | | | | | | |
| Acute | 5.91E-05 | 4.84E-02 | 1.45E-04 | 1.03E-01 | 34 | 26 | 162 | 65 |
| Mild | 1.10E-02 | 8.16E-01 | 1.86E-02 | 8.16E-01 | 34 | 26 | 52 | 28 |
| Moderate | 1.10E-04 | 6.31E-02 | 2.50E-04 | 1.19E-01 | 34 | 26 | 64 | 22 |
| Severe | 8.37E-07 | 1.06E-03 | 2.44E-06 | 4.23E-03 | 34 | 26 | 46 | 15 |
| **Eosinophils** | | | | | | | | |
| Acute | 3.50E-03 | 4.06E-02 | 6.25E-03 | 9.29E-02 | 40 | 34 | 244 | 185 |
| Mild | 2.12E-02 | 1.79E-01 | 3.08E-02 | 2.42E-01 | 40 | 34 | 74 | 57 |
| Moderate | 5.82E-04 | 1.34E-01 | 1.16E-03 | 2.05E-01 | 40 | 34 | 88 | 64 |
| Severe | 4.63E-02 | 1.38E-02 | 6.17E-02 | 4.02E-02 | 40 | 34 | 82 | 64 |
| **Monocytes** | | | | | | | | |
| Acute | 6.08E-01 | 4.92E-01 | 6.48E-01 | 5.25E-01 | 40 | 38 | 259 | 253 |
| Mild | 5.17E-01 | 1.86E-01 | 5.70E-01 | 2.42E-01 | 40 | 38 | 78 | 73 |
| Moderate | 9.14E-01 | 3.15E-01 | 9.14E-01 | 3.60E-01 | 40 | 38 | 96 | 95 |
| Severe | 1.51E-02 | 4.37E-01 | 2.42E-02 | 4.82E-01 | 40 | 38 | 85 | 85 |
| **Neutrophils** | | | | | | | | |
| Acute | 1.51E-09 | 1.52E-04 | 8.06E-09 | 9.75E-04 | 40 | 36 | 259 | 238 |
| Mild | 3.18E-04 | 5.96E-02 | 6.79E-04 | 1.19E-01 | 40 | 36 | 78 | 68 |
| Moderate | 1.00E-08 | 3.72E-03 | 3.57E-08 | 1.32E-02 | 40 | 36 | 96 | 88 |
| Severe | 1.22E-12 | 3.12E-07 | 1.96E-11 | 4.99E-06 | 40 | 36 | 85 | 82 |
| **NK cells** | | | | | | | | |
| Acute | 4.04E-02 | 9.83E-03 | 5.62E-02 | 3.14E-02 | 40 | 23 | 257 | 146 |
| Mild | 6.59E-01 | 1.28E-01 | 6.80E-01 | 2.05E-01 | 40 | 23 | 77 | 47 |
| Moderate | 1.83E-02 | 2.67E-02 | 2.79E-02 | 7.13E-02 | 40 | 23 | 96 | 49 |
| Severe | 3.51E-03 | 4.86E-04 | 6.25E-03 | 2.54E-03 | 40 | 23 | 84 | 50 |
| **T cells** | | | | | | | | |
| Acute | 1.26E-09 | 5.89E-07 | 8.04E-09 | 6.28E-06 | 40 | 38 | 259 | 251 |
| Mild | 4.46E-05 | 5.55E-04 | 1.19E-04 | 2.54E-03 | 40 | 38 | 78 | 72 |
| Moderate | 7.62E-09 | 1.68E-05 | 3.05E-08 | 1.35E-04 | 40 | 38 | 96 | 94 |
| Severe | 3.54E-13 | 8.09E-10 | 1.13E-11 | 2.59E-08 | 40 | 38 | 85 | 85 |

Table S4: Significance of differences in cell-type proportions (Welch's t-test, Benjamini-Hochberg correction) between follow-up samples and controls. Cell-types predicted by the model are compared to manually labelled. Non-zero values refer to number of samples where at least one cell of given cell-type was labelled or predicted. P-values for rows labelled as Acute are calculated between controls and all follow-up samples pooled; for other rows they are between controls and a given severity only. Related to Figure 5.

| | **P-value** | | **Q-value** | | **Non-zero controls** | | **Non-zero follow-up** | |
|---|---|---|---|---|---|---|---|---|
| | Predicted | Labelled | Predicted | Labelled | Predicted | Labelled | Predicted | Labelled |
| | | | | **B cells** | | | | |
| Acute | 1.21E-01 | 2.01E-01 | 3.43E-01 | 8.23E-01 | 39 | 35 | 103 | 92 |
| Mild | 2.10E-01 | 2.33E-01 | 3.43E-01 | 8.23E-01 | 39 | 35 | 27 | 25 |
| Moderate | 1.53E-01 | 2.01E-01 | 3.43E-01 | 8.23E-01 | 39 | 35 | 34 | 31 |
| Severe | 1.07E-01 | 2.83E-01 | 3.43E-01 | 8.23E-01 | 39 | 35 | 42 | 36 |
| | | | | **Basophils** | | | | |
| Acute | 2.38E-01 | 3.73E-01 | 3.63E-01 | 8.23E-01 | 40 | 24 | 102 | 94 |
| Mild | 9.08E-01 | 4.04E-01 | 9.49E-01 | 8.23E-01 | 40 | 24 | 26 | 24 |
| Moderate | 2.14E-01 | 2.05E-01 | 3.43E-01 | 8.23E-01 | 40 | 24 | 34 | 32 |
| Severe | 5.14E-01 | 6.97E-01 | 5.87E-01 | 8.23E-01 | 40 | 24 | 42 | 38 |
| | | | | **DC cells** | | | | |
| Acute | 2.77E-02 | 8.22E-01 | 2.46E-01 | 8.49E-01 | 34 | 26 | 93 | 63 |
| Mild | 2.09E-02 | 6.84E-01 | 2.46E-01 | 8.23E-01 | 34 | 26 | 24 | 19 |
| Moderate | 4.75E-02 | 7.20E-01 | 2.46E-01 | 8.23E-01 | 34 | 26 | 34 | 20 |
| Severe | 8.30E-02 | 6.06E-01 | 3.15E-01 | 8.23E-01 | 34 | 26 | 35 | 24 |
| | | | | **Eosinophils** | | | | |
| Acute | 5.39E-02 | 5.05E-01 | 2.46E-01 | 8.23E-01 | 40 | 34 | 104 | 94 |
| Mild | 4.27E-01 | 9.55E-01 | 5.25E-01 | 9.55E-01 | 40 | 34 | 27 | 25 |
| Moderate | 1.89E-01 | 6.42E-01 | 3.43E-01 | 8.23E-01 | 40 | 34 | 35 | 31 |
| Severe | 3.20E-02 | 3.09E-01 | 2.46E-01 | 8.23E-01 | 40 | 34 | 42 | 38 |
| | | | | **Monocytes** | | | | |
| Acute | 3.18E-01 | 4.49E-01 | 4.42E-01 | 8.23E-01 | 40 | 38 | 103 | 99 |
| Mild | 6.69E-01 | 7.97E-01 | 7.38E-01 | 8.49E-01 | 40 | 38 | 27 | 27 |
| Moderate | 3.47E-01 | 2.71E-01 | 4.44E-01 | 8.23E-01 | 40 | 38 | 34 | 32 |
| Severe | 1.45E-01 | 3.19E-01 | 3.43E-01 | 8.23E-01 | 40 | 38 | 42 | 40 |
| | | | | **Neutrophils** | | | | |
| Acute | 1.98E-01 | 6.71E-01 | 3.43E-01 | 8.23E-01 | 40 | 36 | 104 | 99 |
| Mild | 9.46E-01 | 6.94E-01 | 9.49E-01 | 8.23E-01 | 40 | 36 | 27 | 27 |
| Moderate | 1.96E-01 | 4.87E-01 | 3.43E-01 | 8.23E-01 | 40 | 36 | 35 | 32 |
| Severe | 1.46E-01 | 5.28E-01 | 3.43E-01 | 8.23E-01 | 40 | 36 | 42 | 40 |
| | | | | **NK cells** | | | | |
| Acute | 4.86E-01 | 3.96E-03 | 5.76E-01 | 6.34E-02 | 40 | 23 | 104 | 99 |
| Mild | 1.82E-01 | 4.36E-02 | 3.43E-01 | 4.65E-01 | 40 | 23 | 27 | 27 |
| Moderate | 3.42E-01 | 1.61E-01 | 4.44E-01 | 8.23E-01 | 40 | 23 | 35 | 32 |
| Severe | 9.49E-01 | 4.95E-04 | 9.49E-01 | 1.58E-02 | 40 | 23 | 42 | 40 |
| | | | | **T cells** | | | | |
| Acute | 3.41E-02 | 6.61E-01 | 2.46E-01 | 8.23E-01 | 40 | 38 | 104 | 96 |
| Mild | 8.85E-02 | 4.08E-01 | 3.15E-01 | 8.23E-01 | 40 | 38 | 27 | 27 |
| Moderate | 3.94E-02 | 7.76E-01 | 2.46E-01 | 8.49E-01 | 40 | 38 | 35 | 31 |
| Severe | 2.94E-01 | 5.23E-01 | 4.28E-01 | 8.23E-01 | 40 | 38 | 42 | 38 |

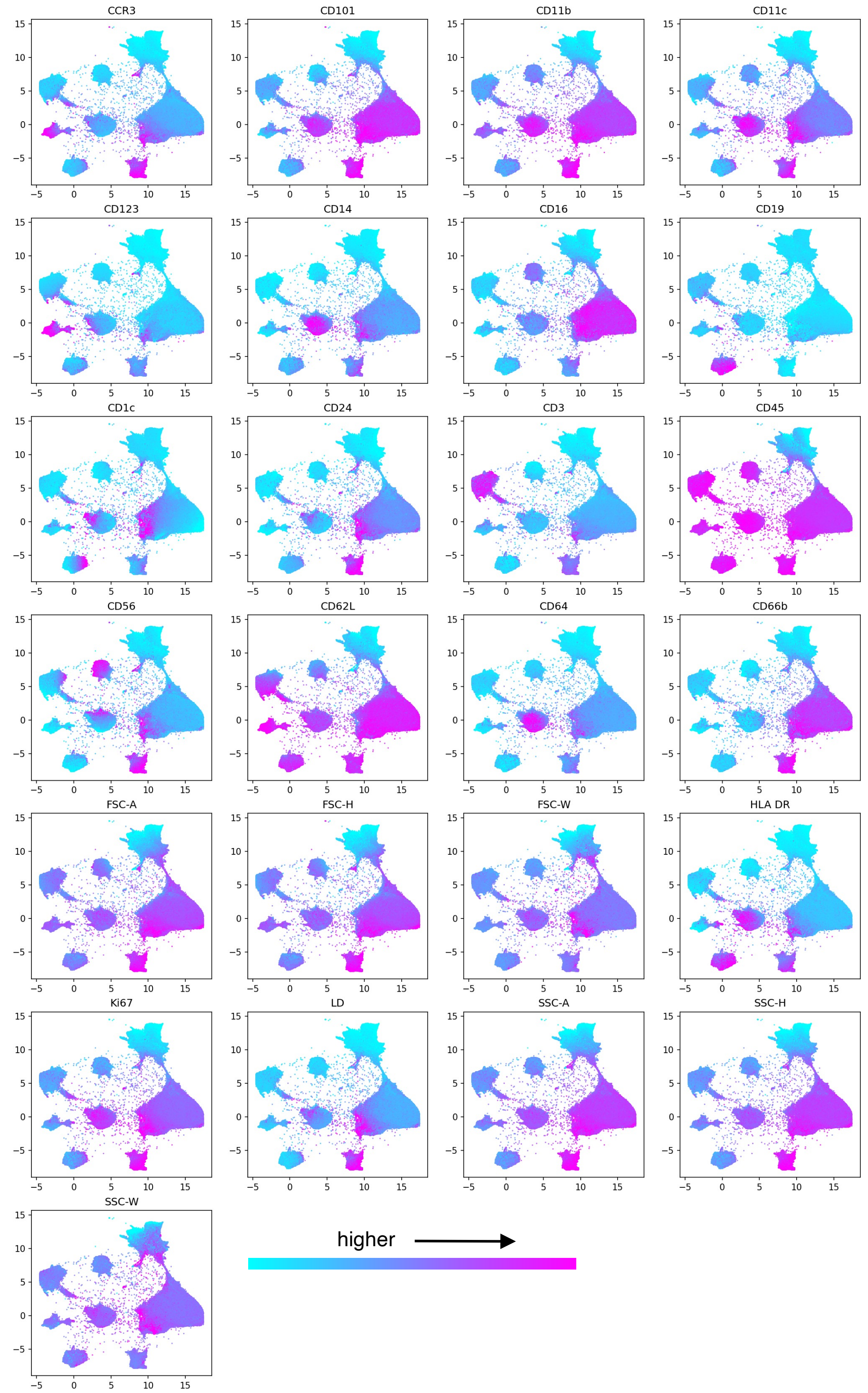


Figure S5: **Marker values in whole blood panels after batch effect correction and imputation**. Related to Figure 4. Integrated latent space including all data samples, shown on 2D UMAP representation. Relative marker values shown within the space.

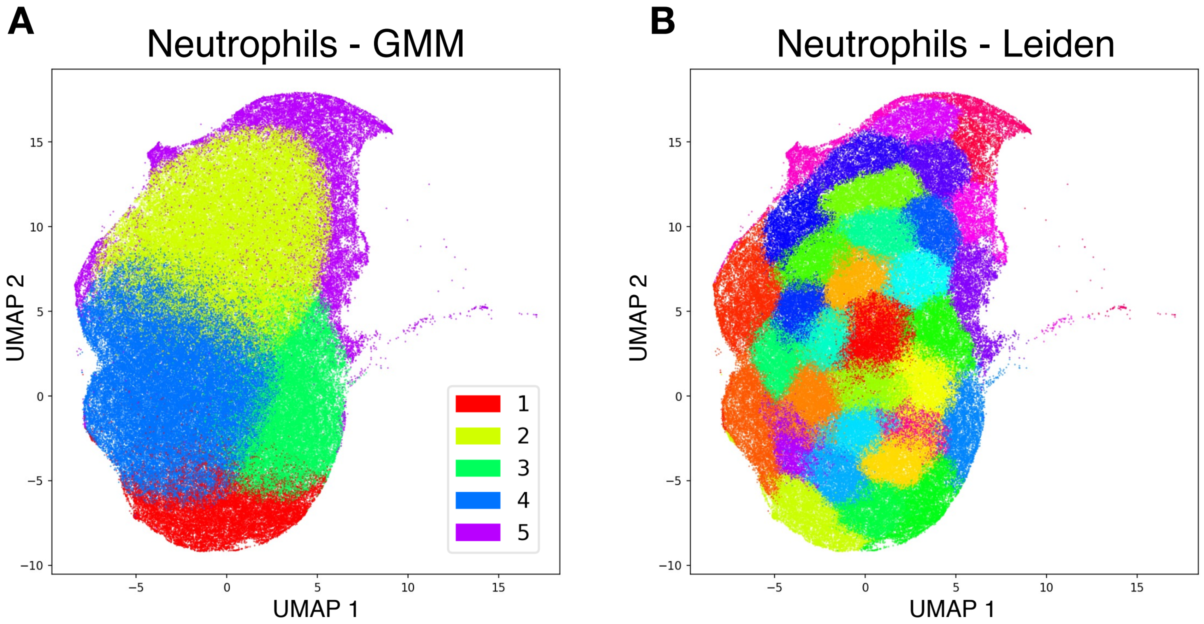


Figure S6: **Clustering of the neutrophil population from the chemokine panel.**. Related to Figure 7. UMAP visualisations of reconstructed data space of chemokine panels after integration, showing clustering of neutrophils. (A) Gaussian Mixture Model clustering (5 clusters specified). (B) Leiden clustering (31 clusters detected). Exact cluster-colour assignment not shown due to space limitation.

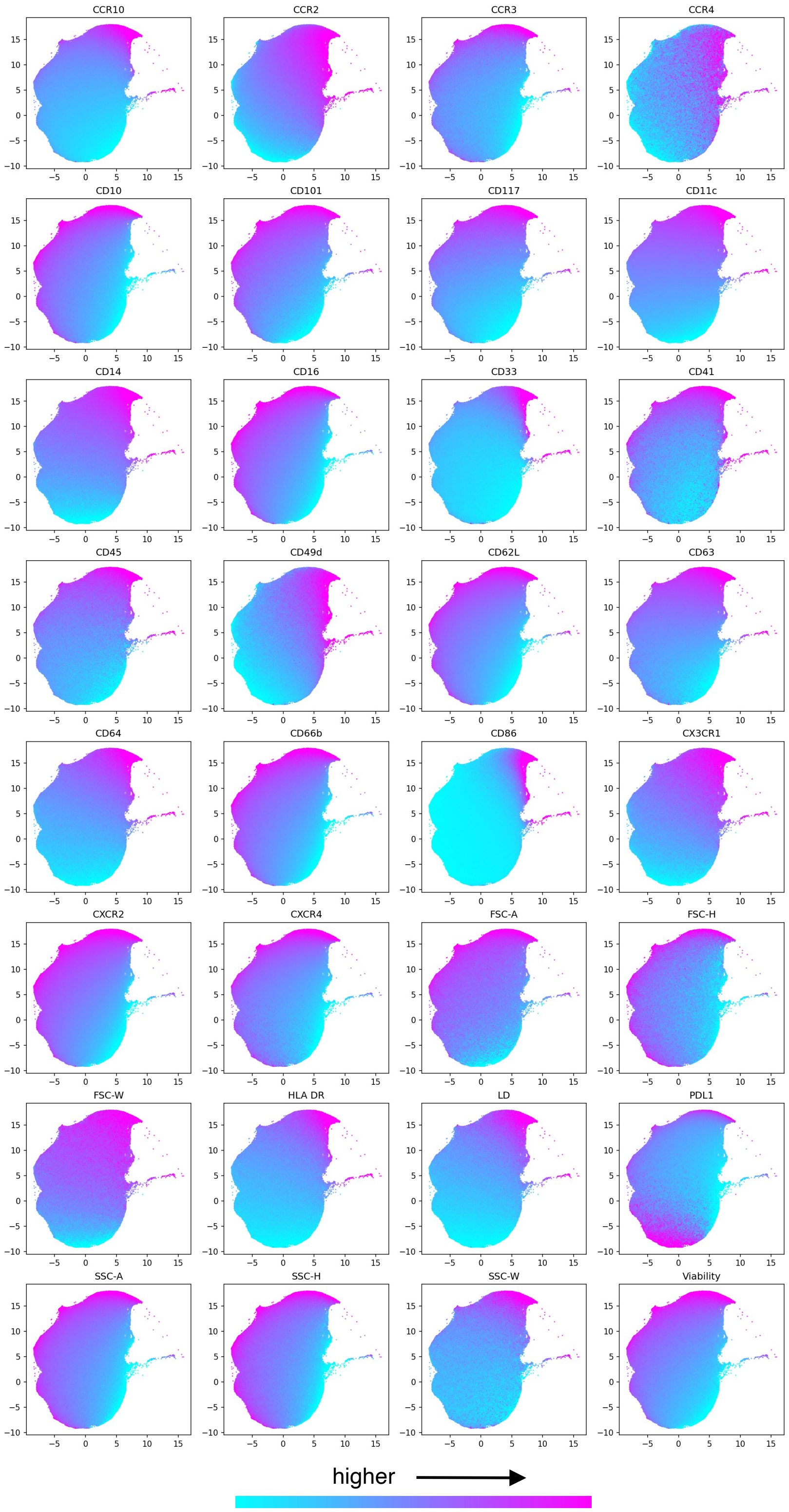


Figure S7: **Marker values of neutrophils in chemokine panels after batch effect correction and imputation**. Related to Figure 6. Integrated latent space including all data samples, shown on 2D UMAP representation. Relative marker values shown within the space.

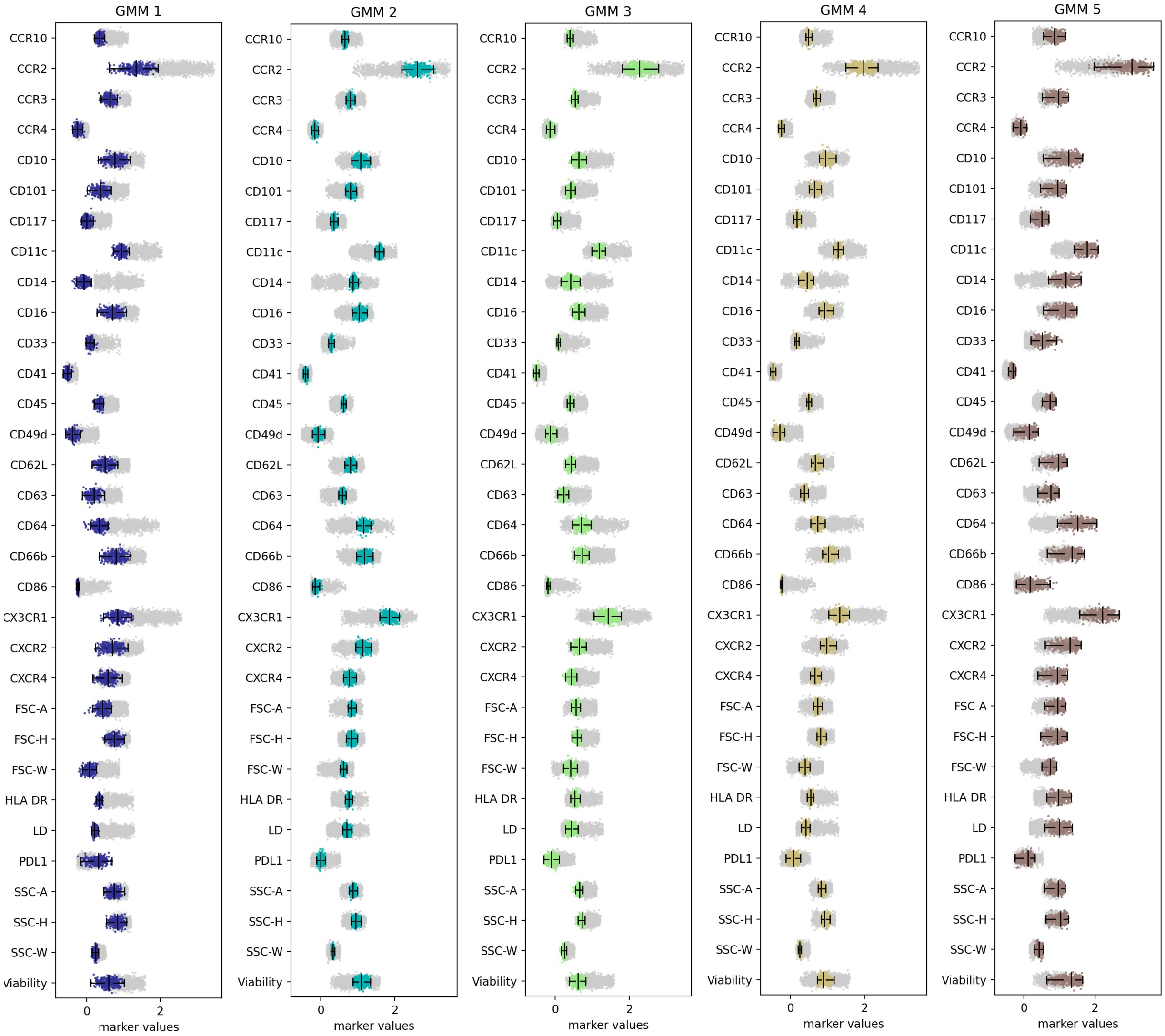


Figure S8: **Relative marker levels for each of neutrophil GMM clusters in imputed values of chemokine panel**. Related to Figure 6 and Figure 7. Vertical lines indicate minimum, median, and maximum mean marker level across all samples, excluding outliers. Coloured values belong to a given cluster, light grey values in the background belong to all other clusters combined.

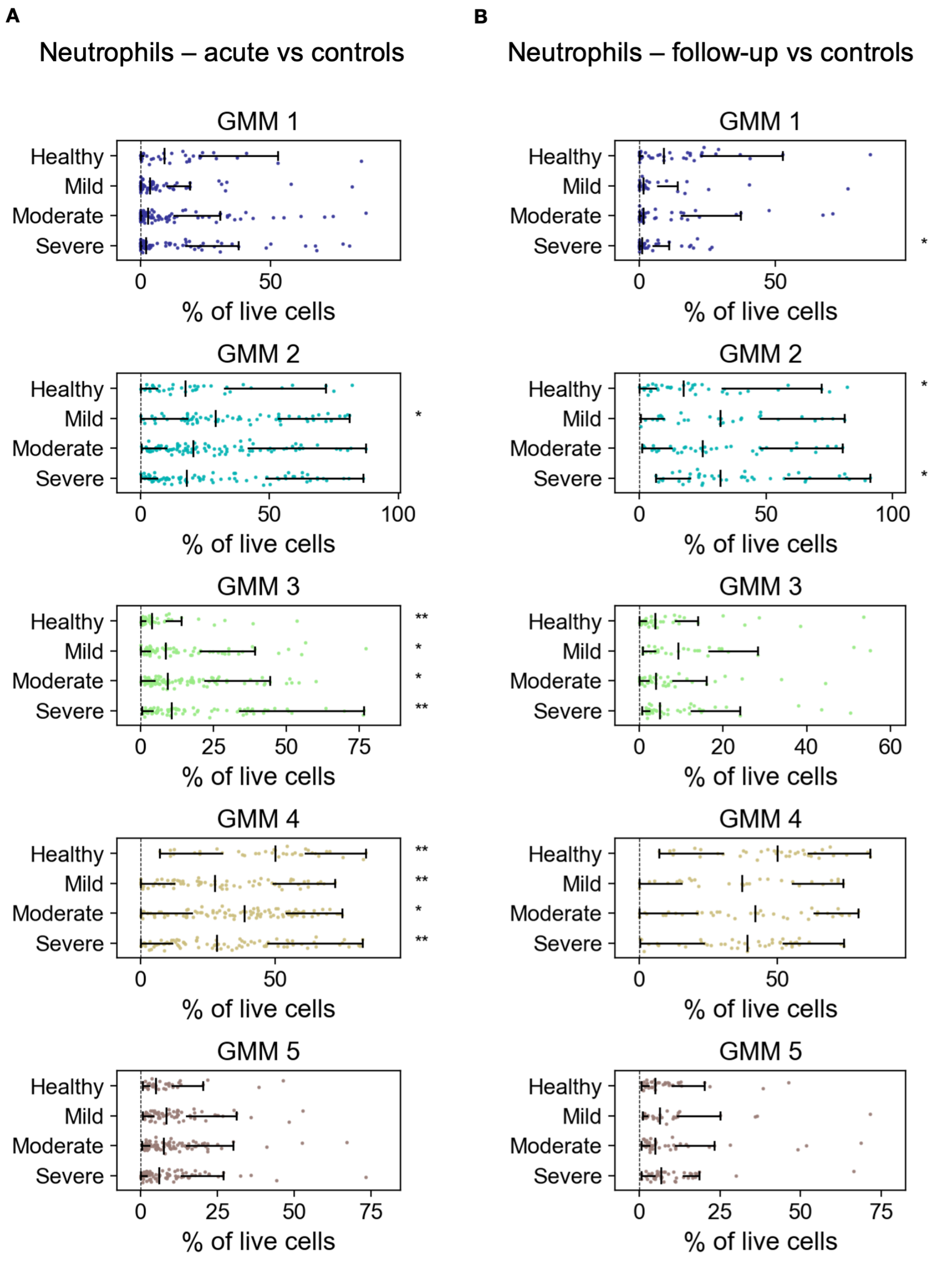


Figure S9: **Distribution of neutrophil GMM cluster proportions across severity groups**. Related to Figure 6 and Figure 7. Vertical lines indicate minimum, median, and maximum proportion, excluding outliers. Significance indicators (*q<0.05, **q<0.01) shown next to each row were obtained from Welch's t-test after Benjamini–Hochberg correction applied within each category (acute or follow-up); stars next to Healthy row indicate significance between healthy controls and combined patient samples of all severities; stars next to other rows indicate significance between particular severity samples and controls. (A) Comparison between controls and acute samples. (B) Comparison between controls and follow-up samples.

Table S5: Significance of differences in marker levels on neutrophils (Welch's t-test) between patient samples and controls. Benjamini–Hochberg correction was applied on an extended set of data where each marker was evaluated for significance using 4 severity groups (pooled acute and 3 severity levels). Values imputed by the model are compared to manually gated values after batch standardisation. P-values are calculated between controls and all types of severity pooled. N/A indicates that insufficient samples were available to calculate significance, either due to missing marker, or neutrophil annotation. Bold values indicate $p<0.05$ or $q<0.05$. Related to Figure 6.

| | **Acute vs. controls** | | | | **Follow-up vs. controls** | | | |
|---|---|---|---|---|---|---|---|---|
| | *P-value* | | *Q-value* | | *P-value* | | *Q-value* | |
| | Imputed | Labelled | Imputed | Labelled | Imputed | Labelled | Imputed | Labelled |
| **CCR10** | 9.49E-01 | 5.43E-01 | 9.92E-01 | 7.82E-01 | 5.09E-02 | N/A | 1.95E-01 | N/A |
| **CCR2** | **5.82E-07** | N/A | **6.40E-05** | N/A | **1.93E-04** | N/A | **1.48E-02** | N/A |
| **CCR3** | **1.99E-02** | 1.39E-01 | 1.05E-01 | 3.38E-01 | 8.29E-01 | 4.63E-01 | 8.82E-01 | 8.51E-01 |
| **CCR4** | **5.30E-05** | 3.91E-01 | **1.27E-03** | 6.50E-01 | **3.87E-04** | N/A | **2.01E-02** | N/A |
| **CD10** | **7.18E-04** | **2.70E-05** | **8.11E-03** | **7.55E-04** | 8.19E-01 | **4.37E-02** | 8.76E-01 | 2.80E-01 |
| **CD101** | **2.48E-02** | **1.29E-06** | 1.29E-01 | **5.69E-05** | 4.31E-01 | **2.40E-03** | 6.35E-01 | 5.91E-02 |
| **CD117** | 6.62E-01 | **1.13E-02** | 9.91E-01 | 5.80E-02 | 6.59E-02 | **2.82E-02** | 2.30E-01 | 2.08E-01 |
| **CD11c** | **3.09E-02** | **8.38E-04** | 1.51E-01 | **1.29E-02** | **4.32E-03** | N/A | **4.61E-02** | N/A |
| **CD14** | **1.94E-02** | 6.68E-02 | 1.03E-01 | 2.00E-01 | **3.58E-03** | 8.55E-01 | **4.52E-02** | 9.83E-01 |
| **CD16** | **1.26E-03** | **1.07E-02** | **1.21E-02** | 5.58E-02 | 9.48E-01 | 2.09E-01 | 9.56E-01 | 6.26E-01 |
| **CD33** | 2.08E-01 | 8.15E-01 | 7.48E-01 | 9.13E-01 | 9.90E-02 | N/A | 3.10E-01 | N/A |
| **CD41** | 7.61E-01 | 7.78E-01 | 9.92E-01 | 9.13E-01 | 1.53E-01 | 5.04E-01 | 3.92E-01 | 8.54E-01 |
| **CD45** | 3.78E-01 | 2.27E-01 | 9.80E-01 | 4.65E-01 | **3.88E-02** | 4.60E-01 | 1.70E-01 | 8.51E-01 |
| **CD49d** | **4.73E-06** | 2.69E-01 | **2.02E-04** | 5.02E-01 | **8.60E-04** | 3.34E-01 | **2.01E-02** | 7.76E-01 |
| **CD62L** | **6.60E-03** | **2.14E-03** | **4.22E-02** | **2.44E-02** | 9.10E-01 | **6.95E-03** | 9.30E-01 | 1.00E-01 |
| **CD63** | 5.09E-01 | 3.46E-01 | 9.81E-01 | 6.09E-01 | **3.93E-02** | 3.38E-01 | 1.70E-01 | 7.76E-01 |
| **CD64** | **7.08E-05** | **1.44E-09** | **1.60E-03** | **4.44E-07** | **5.66E-04** | 1.87E-01 | **2.01E-02** | 5.92E-01 |
| **CD66b** | **4.80E-03** | 9.32E-01 | **3.35E-02** | 9.57E-01 | 8.80E-01 | 3.91E-01 | 9.11E-01 | 8.19E-01 |
| **CD86** | **1.92E-09** | 5.73E-01 | **7.38E-07** | 7.95E-01 | **4.08E-03** | N/A | **4.61E-02** | N/A |
| **CX3CR1** | **4.09E-05** | N/A | **1.10E-03** | N/A | **2.47E-04** | N/A | **1.58E-02** | N/A |
| **CXCR2** | **5.38E-03** | **1.70E-03** | **3.69E-02** | **2.16E-02** | 7.43E-01 | 1.67E-01 | 8.25E-01 | 5.63E-01 |
| **CXCR4** | **1.23E-02** | **7.02E-03** | 7.15E-02 | **4.59E-02** | 9.30E-01 | 1.36E-01 | 9.42E-01 | 5.45E-01 |
| **FSC-A** | 1.55E-01 | **2.82E-02** | 5.85E-01 | 1.14E-01 | 3.88E-01 | **2.39E-02** | 6.08E-01 | 1.90E-01 |
| **FSC-H** | **9.38E-05** | **4.36E-03** | **1.76E-03** | **3.44E-02** | 5.58E-02 | **6.34E-03** | 2.08E-01 | 9.61E-02 |
| **FSC-W** | 9.23E-02 | 9.21E-01 | 4.03E-01 | 9.52E-01 | **5.89E-03** | 5.13E-01 | 5.30E-02 | 8.55E-01 |
| **HLA DR** | **8.96E-05** | 2.69E-01 | **1.76E-03** | 5.02E-01 | **8.94E-04** | N/A | **2.01E-02** | N/A |
| **LD** | **1.56E-02** | 4.58E-01 | 8.54E-02 | 7.04E-01 | **2.15E-03** | 4.78E-01 | **3.60E-02** | 8.51E-01 |
| **PDL1** | **9.34E-03** | 5.80E-01 | 5.60E-02 | 7.95E-01 | **4.53E-02** | 9.36E-01 | 1.81E-01 | 9.91E-01 |
| **PDL2** | | 1.11E-01 | | 2.96E-01 | | 7.19E-02 | | 3.98E-01 |
| **SSC-A** | **4.37E-04** | **3.97E-03** | **5.41E-03** | **3.44E-02** | 4.66E-01 | **6.05E-03** | 6.39E-01 | 9.61E-02 |
| **SSC-H** | **1.19E-04** | **7.66E-03** | **1.98E-03** | **4.59E-02** | 2.07E-01 | **3.34E-02** | 4.49E-01 | 2.26E-01 |
| **SSC-W** | 1.43E-01 | 8.18E-01 | 5.59E-01 | 9.13E-01 | **7.57E-03** | 7.82E-01 | 6.06E-02 | 9.65E-01 |
| **Viability** | **3.39E-02** | N/A | 1.61E-01 | N/A | 5.01E-01 | N/A | 6.53E-01 | N/A |

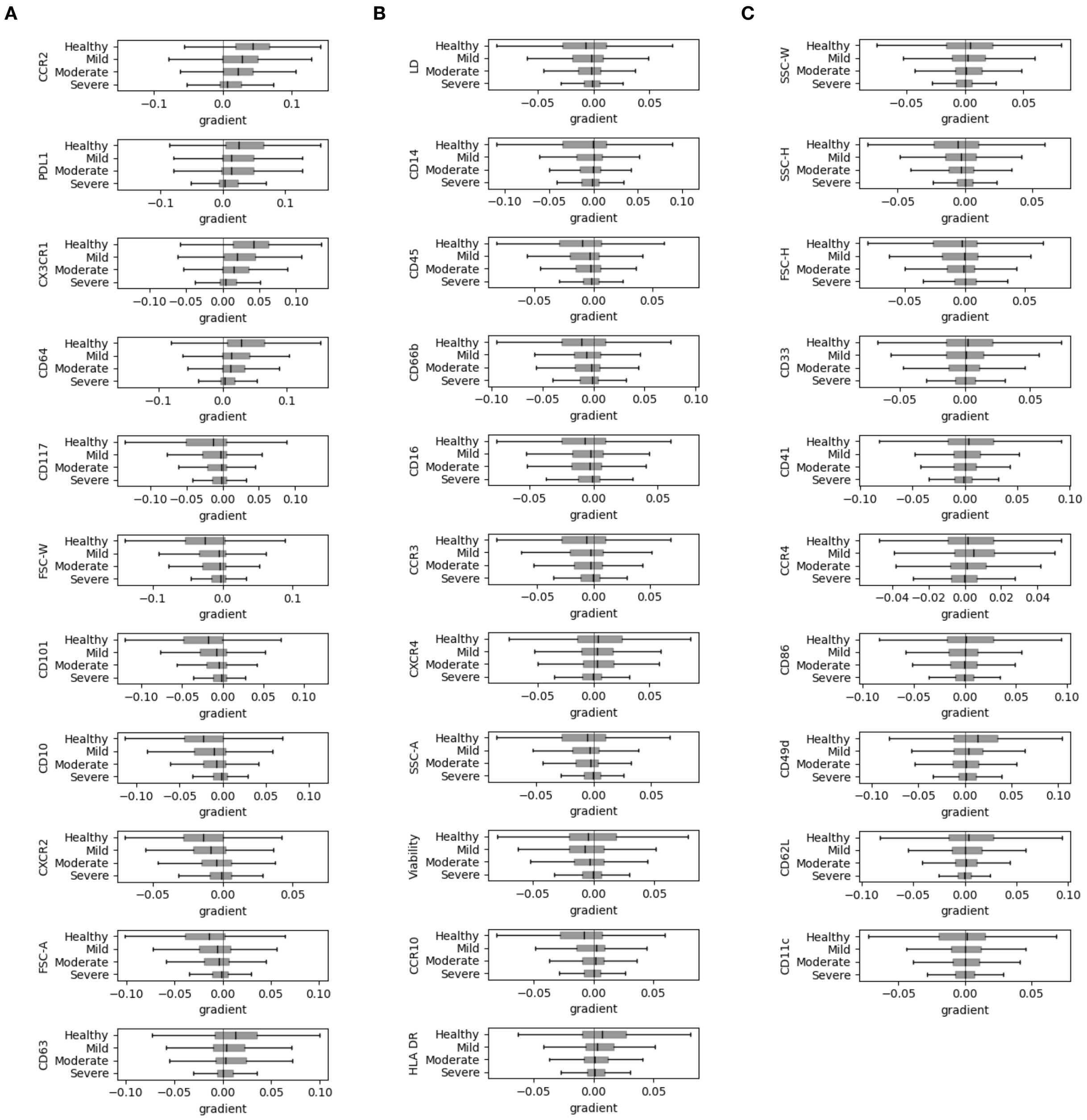


Figure S10: **Neutrophil marker gradients correlated with increase in predicted severity**. Related to Figure 6. Average gradient for each marker on neutrophils input w.r.t. regression output predicting increasing COVID severity. Gradients were obtained from a set model trained using only chemokine panel input, and are grouped by the predicted severity of the sample. (A) Markers sorted from strongest average gradient across all severities to weakest. (B) Continued. (C) Continued.

Table S6: **Default hyper-parameters and optimisation ranges.** Related to STAR Methods: *Model Selection and Hyperparameter Optimisation*.

| Hyper-parameter | default | min | max |
|---|---|---|---|
| latent_dim | 50 | 20 | 100 |
| hidden | 2 | 1 | 2 |
| width | 256 | 16 | 1024 |
| relu_slope | 0.2 | 0 | 0.3 |
| dropout | 0.0 | 0 | 0.3 |
| pull | 1.0 | 0.1 | 10.0 |
| cond_dim | 10 | 10 | 20 |
| cond_hidden | 0 | 0 | 1 |
| cond_width | 256 | 16 | 512 |
| lr_unsupervised | 1.0 | 0.5 | 2.0 |
| lr_supervised | 1.0 | 0.5 | 2.0 |
| lr_merge | 1.0 | 0.5 | 2.0 |
| grad_clip | 0.1 | 0.0001 | 1.0 |
| ease_epochs | 1 | 1 | 9 |
| frequency | 1.0 | 0.1 | 3.0 |
| batch_size | 512 | 128 | 1024 |
| beta | 1.0 | 0.0 | 1.0 |

Table S7: **Training and generation speed.** Related to STAR Methods: *UVAE Framework: General Overview*. Times listed below were evaluated on a M1 Macbook Pro, for a dataset including 100,000 samples (one epoch of training, or when generating). Main model is configured for 3 panels, with conditioning, normalization, subspace merging, MMD and resampling; imputation from 3 panels merged. Imputation model is configured for 3 panels with 3 subspace merging constraints; imputation from a single chosen panel.

| | Main training | Adding additional panel | Generating samples |
|---|---|---|---|
| **Main model (lineage)** | 22s / epoch | 17s / epoch | 3.4s |
| **Imputation model** | 9s / epoch | - | 0.63s |
